# Bayesian Inference for a Generative Model of Transcriptome Profiles from Single-cell RNA Sequencing

**DOI:** 10.1101/292037

**Authors:** Romain Lopez, Jeffrey Regier, Michael Cole, Michael Jordan, Nir Yosef

## Abstract

Transcriptome profiles of individual cells reflect true and often unexplored biological diversity, but are also affected by noise of biological and technical nature. This raises the need to explicitly model the resulting uncertainty and take it into account in any downstream analysis, such as dimensionality reduction, clustering, and differential expression. Here, we introduce Single-cell Variational Inference (scVI), a scalable framework for probabilistic representation and analysis of gene expression in single cells. Our model uses variational inference and stochastic optimization of deep neural networks to approximate the parameters that govern the distribution of expression values of each gene in every cell, using a non-linear mapping between the observations and a low-dimensional latent space.

By doing so, scVI pools information between similar cells or genes while taking nuisance factors of variation such as batch effects and limited sensitivity into account. To evaluate scVI, we conducted a comprehensive comparative analysis to existing methods for distributional modeling and dimensionality reduction, all of which rely on generalized linear models. We first show that scVI scales to over one million cells, whereas competing algorithms can process at most tens of thousands of cells. Next, we show that scVI fits unseen data more closely and can impute missing data more accurately, both indicative of a better generalization capacity. We then utilize scVI to conduct a set of fundamental analysis tasks – including batch correction, visualization, clustering and differential expression – and demonstrate its accuracy in comparison to the state-of-the-art tools in each task. scVI is publicly available, and can be readily used as a principled and inclusive solution for multiple tasks of single-cell RNA sequencing data analysis.

## 1 Introduction

Single-cell RNA sequencing (scRNA-seq) is an increasingly popular tool that opens the way for studying cellular heterogeneity at a high resolution [1], thus shedding new light on fundamental questions in areas such as development [2], autoimmunity [3], and cancer [4]. Interpreting scRNA-seq remains challenging, however, as the data is confounded by nuisance factors such as variation in capture efficiency and sequencing depth [5], amplification bias, batch effects [6] and transcriptional noise [7]. To avoid mistaking nuisance variation for relevant biological diversity, one must therefore account for measurement bias and uncertainty, especially due to the highly abundant false negatives or “dropout” events [8].

The challenge of modeling bias and uncertainty in single-cell data has been explored by several recent studies. A common theme in these studies is treating each data point (cell × gene) as a random variable and fitting a parametric statistical model to this variable. Most existing models are built on a mixture of an “expression” component, which is usually modeled as a negative binomial (e.g., ZINB-WaVE [9]) or log normal (e.g., BISCUIT [10] and ZIFA [11]), and a zero (or low expression) component. The parameters of the model are determined by a combination of cell- and gene-level coefficients, and in some cases additional covariates provided as metadata (e.g., biological condition, batch, and cell quality [9]). All of these methods can therefore be interpreted as finding a low-dimensional representation of the data which can be used to approximate the parameters of the cell × gene random variables. Once these models have been fit to the data, they can then in principle be used for various downstream tasks such as normalization (e.g., scaling, correcting batch effects), imputation of missing data, visualization and clustering.

A complementary line of studies focuses on only one of these tasks, often without explicit probabilistic modeling. For instance, SIMLR [12] fits a cell-cell similarity matrix, under the assumption that this matrix has a block structure with a fixed number of clusters. The resulting model can be used for clustering and for visualization [13]. MAGIC [14] performs imputation of unobserved (dropout) counts by propagation in a cell-cell similarity graph. Census [15] and SCNorm [16] look for proper scaling factors by explicitly modeling the dependence of gene expression on sequencing depth or spike-in RNA. For differential expression analysis, the most common methods consist of both methods developed for bulk count data (e.g., DESeq2 [17] and edgeR [18]) as well as methods developed for scRNA-seq data, explicitly accounting for the high dropout rates (e.g., MAST [19]).

While these methods yield insights into biological variation in single-cell data, several significant limitations remain. First, all of the existing distributional modeling methods assume that a low-dimensional manifold underlies the data, and that the mapping from this manifold to the parameters of the model can be captured by a generalized linear model. While the notion of a restricted dimensionality is plausible (reflecting, e.g., common regulatory mechanisms among genes or common states among cells), it is difficult to justify the assumption of linearity. Second, different existing methods use their fitted models for different subsets of tasks (e.g., imputation and clustering, but not differential expression [10]). Ideally, one would have a single distributional model that would be used for a range of downstream tasks, thus help ensuring consistency and interpretability of the results. Finally, computational scalability is increasingly important. While most existing methods can be applied to no more than tens of thousands of cells, the next generation of tools must scale to the size of recent data sets (commercial [20], or envisioned by consortia such as the Human Cell Atlas [21]) that consist of hundreds of thousands of cells or more.

To address these limitations, we developed a fully probabilistic approach to normalization and downstream analysis of scRNA-seq data, which we refer to as Single-cell Variational Inference (scVI). scVI is based on a hierarchical Bayesian model [22] with conditional distributions specified by deep neural networks. The transcriptome of each cell is encoded through a non-linear transformation into a low-dimensional latent vector of normal random variables. This latent representation is then decoded by another non-linear transformation to generate a posterior estimate of the distributional parameters of each gene in each cell, assuming a zero-inflated negative binomial distribution - a commonly accepted distributional model for gene expression count data that accounts for the observed over-dispersion and limited sensitivity [23, 17, 9]. Notably, recent work [24] shows how neural networks can be used as useful function approximators for single-cell RNA sequencing data; however this work does not use distributional modeling, and thus limited in its scope and applicability to downstream tasks.

In the remainder of this paper, we demonstrate the extent to which scVI addresses the current methodological limitations. First, we demonstrate the scalability of scVI to data sets of up to a million cells. Second, we show that, by using non-linear transformations, scVI better fits unseen data (imputation and held-out log-likelihood). Finally, we demonstrate that the model of scVI can be used for a number of tasks, including batch removal and normalization, clustering, dimensionality reduction and visualization, and differential expression. For each of these tasks, we show that scVI compares favorably to the current state-of-the-art methods. The implementation of scVI is based on the TensorFlow library [25], and is publicly available at https://github.com/YosefLab/scVI.

## 2 Results

### 2.1 Model definition

The primary output of a scRNA-seq experiment is an *N × G*-matrix *x* that records the number of transcripts measured for each of *G* genes in each of *N* cells. We may also have a batch annotation *s_n_* observed for each cell *n* as well.

We model the expression level *x_ng_* measured for each cell *n* and gene *g* as a sample drawn from a conditional distribution that has a zero-inflated negative binomial (ZINB) form [23, 17, 9]. The distribution is conditioned on the observed batch annotation, as well as two additional, unobserved random variables. The latent random variable *ℓ_n_* represents nuisance variation due to variation in capture efficiency and sequencing depth. It is drawn from a log-normal distribution and serves as a cell-specific scaling factor.

The latent random variable *z_n_* represents the remaining variability, which should better reflect biological differences between cells. It is drawn from a standard multivariate normal of low dimensionality *d*, and provides a latent-space representation that can be used for visualization and clustering. The reason for drawing *z_n_* from a multivariate normal is essentially for computational convenience (see Methods 4.1). The matrix *ρ* is an intermediate value that relates the observations *x_ng_* to the latent variables. It provides a batch-corrected, normalized estimate of the percentage of transcripts in each cell *n* that originate from each gene *g*. We use *ρ* for differential expression analysis, and its scaled version (multiplying by the estimated library size) for imputation.

Altogether, each expression value *x_ng_* is drawn independently through the following process:

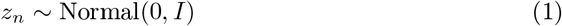

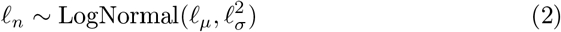

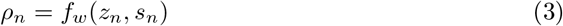

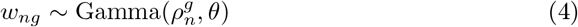

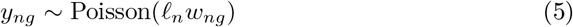

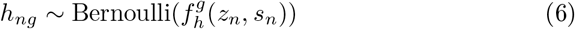

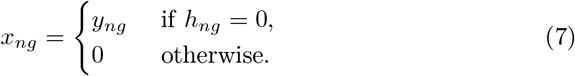

Here *B* denotes the number of batches and 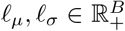 parameterize the prior for the scaling factor (on a log scale). The specification of these parameters is discussed in Methods 4.1. The parameter 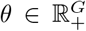 denotes a gene-specific inverse dispersion, estimated via variational Bayesian inference (Methods 4.2). *f_w_* and *f_h_* are neural networks that map the latent space and batch annotation back to the full dimension of all genes: ℝ^*d*^ × {0, 1}*^B^ →*ℝ ^*G*^ (Figure 1b, NN5-6). We use superscript annotation (e.g., 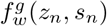 to refer to a single entry that corresponds to a specific gene *g*. We enforce 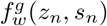 to take values in the probability simplex (namely for each cell *n* the sum of 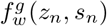 values over all genes *g* is one), thus providing interpretation as expected frequencies. Importantly, neural networks allows us to go beyond the generalized linear model framework and provide a more flexible model of gene expression. Figure 1a specifies the complete graphical model and its implementation using neural-network conditionals. Methods 4.1 provides further details on the specification of this probabilistic model.

**Figure 1:**
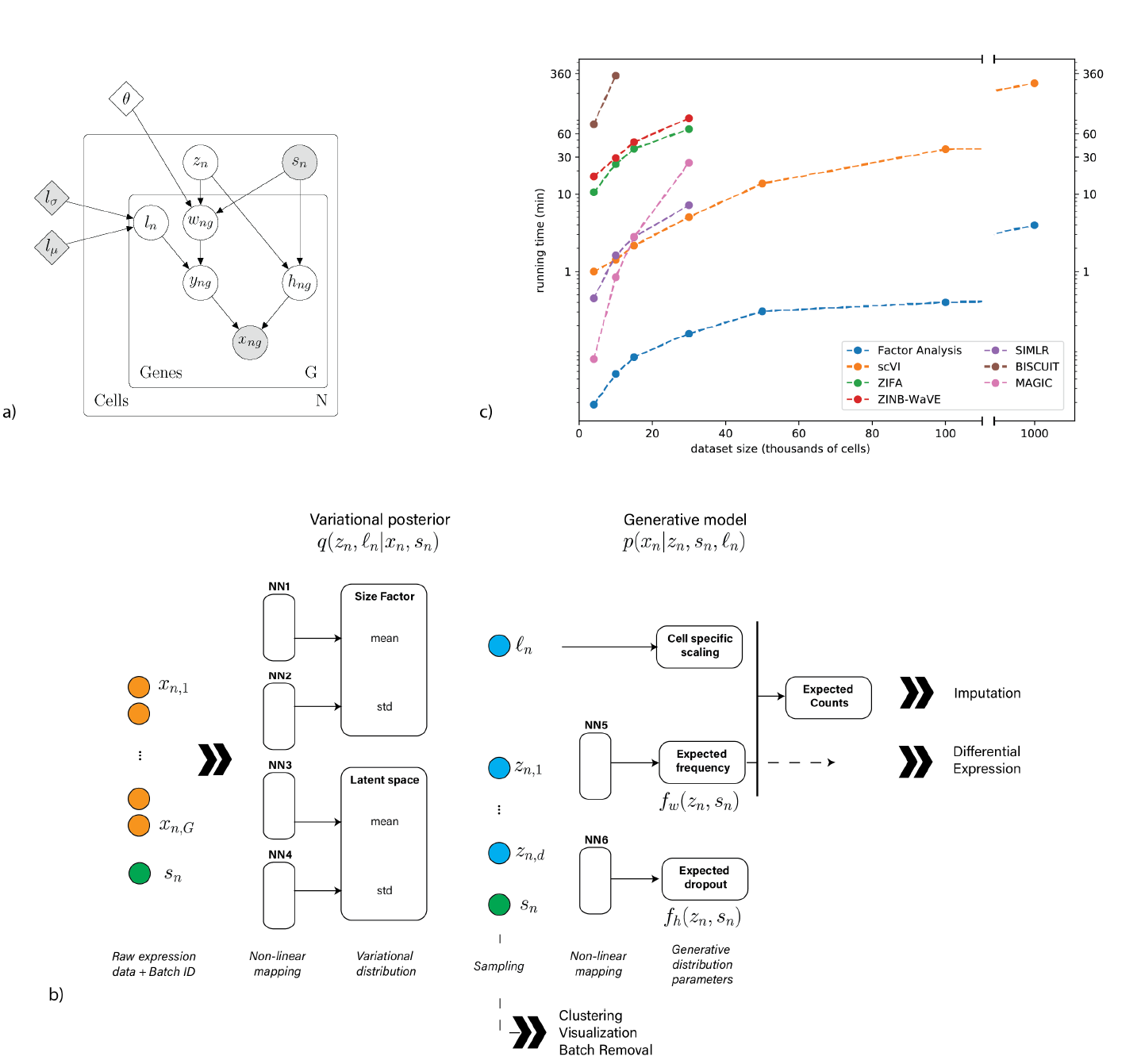
Overview of scVI. Given a gene-expression matrix with batch annotations as input, scVI learns a non-linear embedding of the cells that can be used for multiple analysis tasks. (a) The underlying graphical model. Shaded vertices represent observed random variables. Empty vertices represent latent random variables. Shaded diamonds represent constants, set a priori. Empty diamonds represent global variables shared across all genes and cells. Edges signify conditional dependency. Rectangles (“plates”) represent independent replication. (b) The computational trees (neural networks) used to compute the embedding as well as the distribution of gene expression. (c) Comparison of running times (y-axis) on the BRAIN-LARGE data with a limited set of 720 genes, and with increasing input sizes (x-axis; cells in each input set are sampled randomly from the complete dataset). scVI is compared against existing methods for dimensionality reduction in the scRNA-seq literature. As control, we also add basic matrix factorization with factor analysis (FA).

The distribution *p*(*x_ng_|z_n_, s_n_, ℓ_n_*) is zero-inflated negative binomial (ZINB) [23] with mean 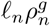, gene-specific dispersion *θ^g^* and zero-inflation probability 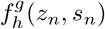 (see Appendix A). Because the marginal distribution *p*(*x_ng_|z_n_, s_n_, ℓ_n_*) is not amenable to exact Bayesian computation, we use variational inference [26] to approximate the posterior distribution. Our variational distribution, *q*(*z_n_*, log *ℓ_n_|x_n_, s_n_*), is Gaussian with a diagonal covariance matrix. The mean and covariance of the variational distribution are given by an encoder network applied to *x_n_* and *s_n_* [27] (Figure 1b, NN1-4). With this formulation, the approximate inference problem can be efficiently solved using a stochastic optimization procedure where we optimize the variational lower bound (Methods 4.2).

### 2.2 Datasets

We apply scVI to seven publicly available datasets (see Methods 4.5 for pre-processing information). We focus on datasets with unique molecular identifiers (UMIs), which prevents overcounting due to amplification. Due to scalability issues, not all the benchmark methods included in this paper are applicable to all datasets. We therefore provide the list of methods applicable to each dataset, along with additional information such as the hyperparameters for scVI, in the Methods section and Supplementary Table 2.

The first dataset (BRAIN-LARGE) consists of 1.3 million mouse brain cells, spanning the cortex, hippocampus and subventricular zone, and profiled with 10x chromium [20]. We use this dataset to demonstrate the scalability of scVI.

The second dataset (CORTEX) consists of 3,005 mouse cortex cells profiled with the Smart-seq2 protocol, with the addition of UMI [28]. To facilitate comparison with other methods, we use a filtered set of 558 highly variable genes as in [10]. The CORTEX dataset exhibits a clear high-level subpopulation structure, which has been inferred by the authors of the original publication using computational tools and annotated by inspection of specific genes or transcriptional programs. Similar levels of annotation are provided with the third and fourth datasets.

The third dataset (PBMC) consists of 12,039 human peripheral blood mononuclear cells profiled with 10x [29].

The fourth dataset (RETINA) includes 27,499 mouse retinal bipolar neurons, profiled in two batches using the Drop-Seq technology [30]. The original annotations of these datasets were used to benchmark scRNA-seq algorithms in several subsequent studies (e.g., [12, 10]).

The fifth dataset (HEMATO) includes 4,016 cells from two batches that were profiled using in-drop. This data provides a snapshot of hematopoietic progenitor cells differentiating into various lineages. We use this dataset as an example for cases where gene expression varies in a continuous fashion (along pseudo-temporal axes) rather than forming discrete subpopulations [31].

The sixth dataset (CBMC) includes 8,617 cord blood mononuclear cells profiled using 10x along with, for each cell, 13 well-characterized mononuclear antibodies [32]. We used this dataset to analyze how the latent spaces inferred by dimensionality-reduction algorithms summarize protein marker abundance.

The seventh dataset (BRAIN-SMALL) consists of 9,128 mouse brain cells profiled using 10x [20]. This dataset is used as a complement to PBMC for our study of zero abundance and quality control metrics correlation with our generative posterior parameters.

### 2.3 Scalability to large datasets

We start by comparing the scalability of scVI to that of state-of-the-art algorithms for imputation and dimensionality reduction (Figure 1c). We evaluate scalability in terms of runtime and memory requirements for increasing numbers of cells, sampled from the complete BRAIN-LARGE dataset. To facilitate comparison to less scalable methods, we limited the analysis to the 720 genes with largest standard deviation across all cells. All the algorithms were tested on a machine with one eight-core Intel i7-6820HQ CPU addressing 32 GB RAM, and one NVIDIA Tesla K80 (GK210GL) GPU addressing 24 GB RAM.

Available memory (RAM) limits scalability of many existing algorithms. Under the hardware and input settings above, we find that BISCUIT runs out of memory when provided with more than 15K cells. MAGIC, ZIFA, SIMLR and ZINB-WaVE can process up to 50K cells before running out of memory. One explanation for this is the explicit storage in memory of the full-data matrix or its derivative (e.g., the cell-cell distance matrix or a proxy whose memory complexity is linear in the number of data points, as in SIMLR and MAGIC).

Focusing on the memory-feasible dataset sizes, we also observe a range of runtimes. For instance, ZIFA, ZINB-WaVE and BISCUIT have a relatively high runtime requirements possibly because their optimization algorithms need to go through all the training data at each each step: the runtime of each iteration scales linearly in the number of samples and linearly in the number of genes.

scVI relies instead on stochastic optimization, sampling a fixed number of cells at each iteration (Methods 4.2). Its time and space complexity per iteration therefore depend only on the number of genes, ensuring scalability both in terms of memory use and processing time. In practice, the algorithm always converged after 250 epochs. In five hours, scVI processed one million cells for a benchmark set of 720 genes. In ten hours, scVI processed one million cells for 10,000 genes.

### 2.4 Goodness of fit and generalization to held-out data

To evaluate the extent to which the different models fit the data, we use a goodness-of-fit score on unseen data, defined by the marginal log-likelihood of a held-out dataset (Methods 4.7). We first partition our data into “training” and “testing” sets and apply the various methods to learn a ten-dimensional latent space and a mapping from this space to the original dimension of the data. We then measure the marginal likelihood *p*(*x*) of the held-out data for the trained model. Exploring a range of training dataset sizes from a few thousand cells to a hundred thousand cells, sampled from the BRAIN-LARGE dataset (Table 1) with 720 genes, we observe that scVI provides the most likely model for the held-out data (consisting of 10K randomly sampled cells that are not in the training set) and that its added accuracy grows as the training dataset size grows. Using the smaller CORTEX dataset for the same analysis—where we partitioned the data as 60% training and 30% testing—yields similar results (Supplementary Figure 9a). For both datasets, scVI and ZINB-WAVE are more accurate than ZIFA or a standard Factor Analysis (FA), thus corroborating that scRNA-seq data is better approximated by a ZINB than a log-normal or a zero-inflated-log-normal.

**Table 1:** Marginal log likelihood for a held-out subset of the brain-cell dataset. NA means we could not run the given algorithm for this sample size. FA denotes Factor Analysis.

| cells | 4K | 10K | 15K | 50K | 100K |
| --- | --- | --- | --- | --- | --- |
| FA | -1175.36 | -1177.35 | -1177.27 | -1171.93 | -1169.86 |
| ZIFA | -1250.44 | -1250.77 | -1250.59 | NA | NA |
| ZINB-WaVE | -1166.39 | -1163.91 | -1163.39 | NA | NA |
| scVI | **-1150.96** | **-1146.59** | **-1144.88** | **-1136.57** | **-1133.94** |

**Table 2:** Presentation of the different datasets, their gene filtering and applicability of algorithms. “_” indicates combinations of algorithm and dataset that were included in this study. “x” indicates that the algorithm took more than four hours to run (ZIFA) or that the computer ran out of memory (others). “NA” indicates datasets where pre-annotated subpopulations were not available, which makes them less useful for application with SIMLR and benchmark of clustering. “NC” indicates the remaining combinations of algorithm and dataset that were not considered for this study. For instance, BRAIN-SMALL was only used to study the correlation of scVI zero probabilities and quality parameters (Figure 6).

|  | dataset properties |  | scVI hyperparameters |  |  | scalability to other algorithms |  |  |  |  |
| --- | --- | --- | --- | --- | --- | --- | --- | --- | --- | --- |
|  | cells | genes | learning rate | layers | neurons | FA | ZIFA | ZINB WaVE | MAGIC | SIMLR |
| **CORTEX** | 3,005 | 558 | 0.0004 | 1 | 128 | — | — | — | — | — |
| **PBMC** | 12,039 | 3,346 | 0.0004 | 1 | 128 | — | x | x | — | — |
| **HEMATO** | 4,016 | 7,397 | 0.0004 | 1 | 128 | — | x | x | — | NC |
| **HEMATO\*** | 4,016 | 700 | 0.0004 | 1 | 128 | — | x | x | — | NC |
| **BRAIN LARGE** | 10,000 | 720 | 0.001 | 1 | 128 | — | — | — | — | NA |
| **BRAIN LARGE** | 15,000 | 720 | 0.001 | 2 | 128 | — | — | — | NC | NA |
| **BRAIN LARGE** | 50,000 | 720 | 0.001 | 3 | 256 | — | x | x | NC | NA |
| **BRAIN LARGE** | 1M | 720 | 0.001 | 3 | 256 | — | x | x | x | NA |
| **BRAIN LARGE** | 1M | 10,000 | 0.001 | 3 | 256 | — | x | x | x | NA |
| **RETINA** | 27,499 | 13,166 | 0.0005 | 1 | 128 | — | x | x | x | — |
| **CBMC** | 8,617 | 600 | 0.0005 | 1 | 128 | — | — | — | NC | — |
| **BRAIN SMALL** | 9,128 | 3,000 | 0.001 | 1 | 128 | NC | NC | NC | NC | NC |

The held-out marginal likelihood becomes a less informative metric when the data is dominated by zero entries (which was not the case for the two datasets reported above because of gene filtering). When zero entries dominate, this test reduces to comparing which algorithm generates a predominance of values close to zero. We therefore turn to imputation benchmarking as a proxy to evaluate the model’s fit on the remaining datasets.

The ability to impute missing values is useful in practical applications in addition to providing an assay for generalization performance [14]. In the following analysis, we benchmark scVI against BISCUIT, ZINB-WaVE and ZIFA, as well as MAGIC, which provides imputation without explicit statistical modeling. To evaluate these methods on a given dataset, we generated a corrupted training set by setting 9% uniformly chosen non-zero entries to zero. We then fit the perturbed dataset with each of the benchmark methods and evaluate them by comparing the imputed values to the original ones (Methods 4.7). Overall, we observe that the imputation accuracy of scVI is higher or comparable (less than one transcript for median error) across all datasets (Figure 2d-f and Supplementary Figure 9b-d).

**Figure 2:**
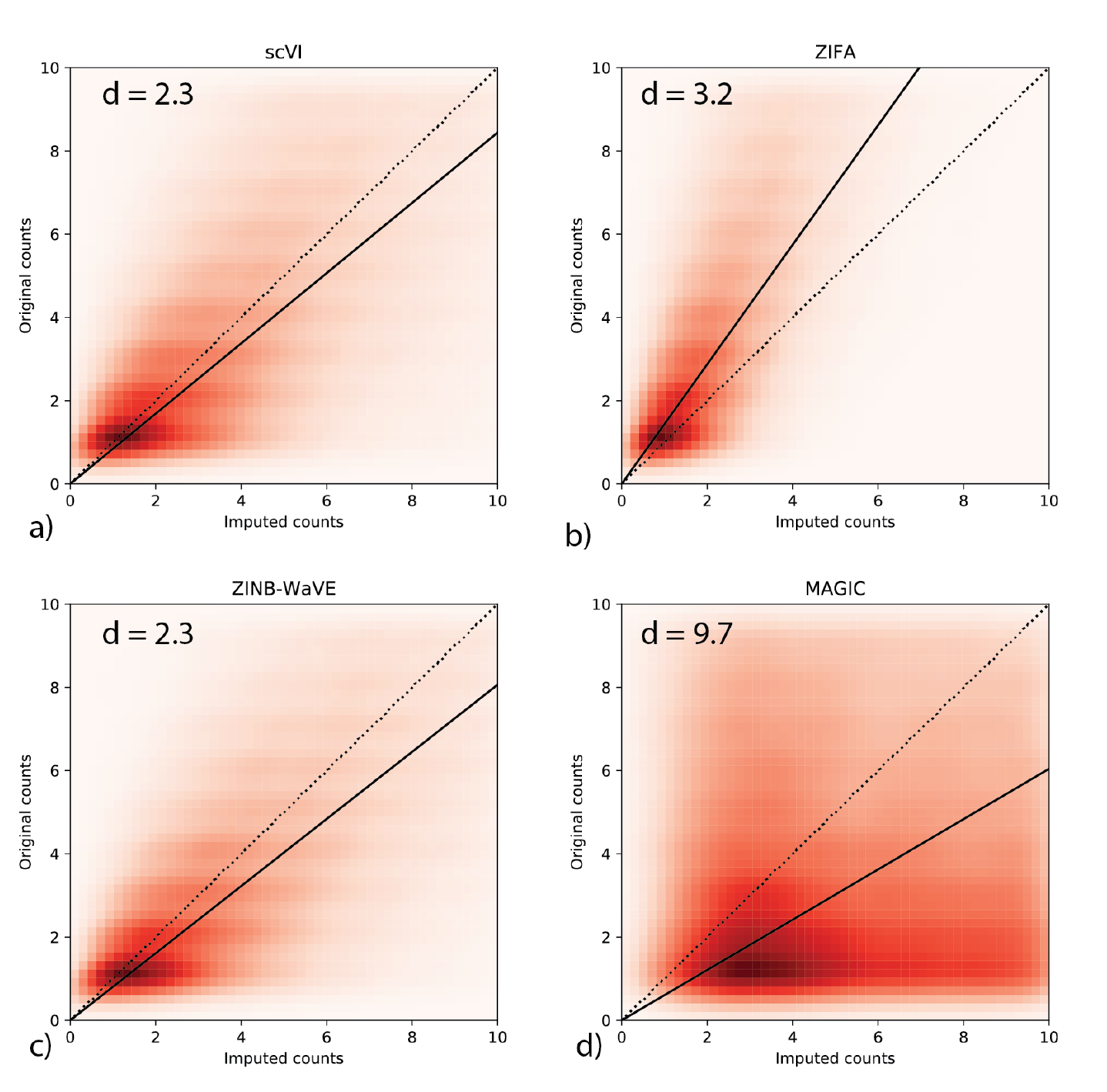
Imputation of scVI on the CORTEX dataset. The heatmaps denote density plots of imputed values (by scVI, ZIFA, MAGIC and ZINB-WaVE respectively) on a down-sampled version versus the original (non-zero) values prior to down-sampling. The reported score *d* is the median imputation error across all the hidden entries (Lower is better; see Methods).

One important exception is the full HEMATO dataset, in which the number of cells (4,016) is smaller than the number of genes (7,397). In such cases, scVI is expected to underfit the data, potentially leading to worse imputation performance. However, additional gene filtering (to the top 700 variable genes) helps to recover an accurate imputation (Supplementary Figure 9d).

To provide an example when the perturbed values depend on the amount of mRNA observed, we also generated a corrupted training set by downsampling 10% uniformly chosen non-zero entries with a binomial law of rate 20%. These values guarantee that most of the dataset is not changed and require the model enough flexibility to impute correctly the changed values. With respect to these corruption scheme, scVI also performs well (Supplementary Figure 11, Supplementary Figure 10).

scVI, like ZIFA and FA, can also be used to generate unseen data by sampling from the latent space. As evidence for the validity of this procedure, we sampled from the posterior of the training data and observed that the resulting “simulated” data is largely consistent with the observed values (Supplementary Figure 7).

### 2.5 Capturing biological structure in a latent space

To further assess the performance of scVI, we evaluated how well its latent space summarizes biological information and recovers biologically coherent subpopulations. For these experiments, we used three datasets where pre-annotated clusters or subpopulations are available: CORTEX, PBMC and RETINA. We then examined whether the annotated subpopulations are distinguishable in the latent space, as in [12]. We report two different metrics for this analysis. First, silhouette width [33], which evaluates whether cells from the same subpopulation have a similar latent representation and cells from different subpopulations have a different representation. Second, we use the latent representation as an input to the *K*-means algorithm, and measure the overlap between the resulting clustering annotations and the pre-specified subpopulations using the Adjusted Rand Index (ARI) and Normalized Mutual Information (NMI) scores (Methods 4.7). For ease of comparison across methods, we set *K* to the number of annotated subpopulations.

While these annotated subpopulations were subject to manual inspection and interpretation, a remaining caveat is that they are computationally derived. To address this we make use of the CBMC dataset that includes measurements of thirteen key marker proteins in addition to mRNA. For evaluation, we quantify how much the similarity between cells in the mRNA latent space resembles their similarity at the protein level. To this end, we compute the overlap fold enrichment between the protein and mRNA-based cell 100-nearest neighbor graph and the Spearman correlation of the adjacency matrices (Methods 4.7).

Based on these benchmarks, we compared scVI to other methods that aim to infer a biologically meaningful latent space (ZIFA, ZINB-WaVE, and FA), using the same clustering scheme. We find that scVI compares favorably to these methods for all the datasets (Supplementary Figure 8ab). Next, we benchmark scVI with SIMLR [12], a method that couples clustering with learning of a cell-cell similarity matrix. For the first set of tests, we set the number of clusters in SIMLR to be the true number of annotated subpopulations. For the CBMC case, we let SIMLR automatically determine that number. When looking at the evaluations that were based on the computationally derived annotations, we find that SIMLR outperforms scVI (Supplementary Figure 8ab). However, while the latent space inferred by SIMLR provides a tight representation for these subpopulations, it may disregard other forms of critical information. Indeed, in the CBMC-based test where the clustering is based on “external” but biologically meaningful data, scVI is the best performing method, albeit by a small margin (Supplemental Figure 8c).

Another example for important information that may be missed is that of a hierarchical structure between clusters, such as the one reported for the CORTEX dataset [28]. We take several cuts at different depths of the hierarchical clustering (Table 3) and report clustering scores based on these agglomerated labels (Supplemental Figure 8efg). These results suggest that scVI and ZINB-WaVE find low-dimensional representations that better preserve this important biological structure.

**Table 3:** Cell-types present in the CORTEX dataset and their labels in some slices of the original hierarchical clustering.

| Cell Type | # cells | Cluster ID<br>(original) | Cluster ID<br>(#4) | Cluster ID<br>(#3) | Cluster ID<br>(#2) |
| --- | --- | --- | --- | --- | --- |
| astrocytes_ependymal | 224 | 0 | 1 | 0 | 0 |
| endothelial-mural | 235 | 1 | 2 | 1 | 0 |
| interneurons | 290 | 2 | 3 | 2 | 1 |
| microglia | 98 | 3 | 2 | 1 | 0 |
| oligodendrocytes | 820 | 4 | 0 | 0 | 0 |
| pyramidal CA1 | 939 | 5 | 3 | 2 | 1 |
| pyramidal SS | 399 | 6 | 3 | 2 | 1 |

**Table 4:**
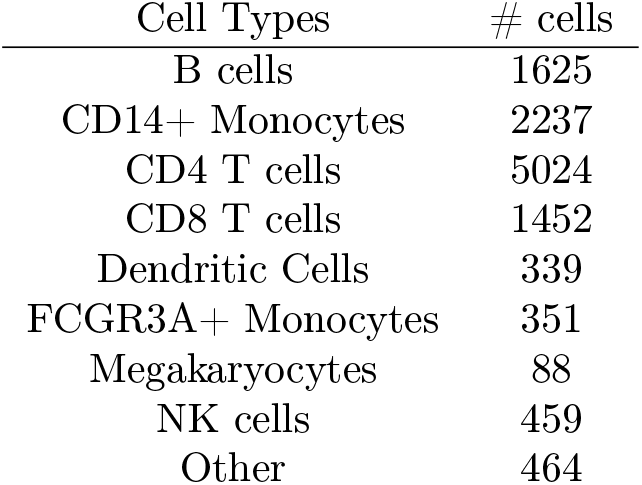
Cell-types present in the PBMC dataset

A second important case is when the variation between cells has a continuous, rather than discrete form. An example for that is the HEMATO dataset, which consists of hematopoietic cells annotated along seven different stages of differentiation. As a first step, we focus on differentiation towards either granulocytic neutrophil or erythoid fate [31]. SIMLR applied to this dataset predicts the presence of five clusters, and the resulting five-nearest-neighbors graph (visualized using a Fruchterman-Reingold force-directed algorithm, see Methods 4.6) does not reflect the continuous nature of this system. Conversely, standard PCA analysis and scVI are able to capture this property of the data (Figure 3), albeit with less precision than a manually tuned process used in the original publication (Supplemental Figure 16).

**Figure 3:**
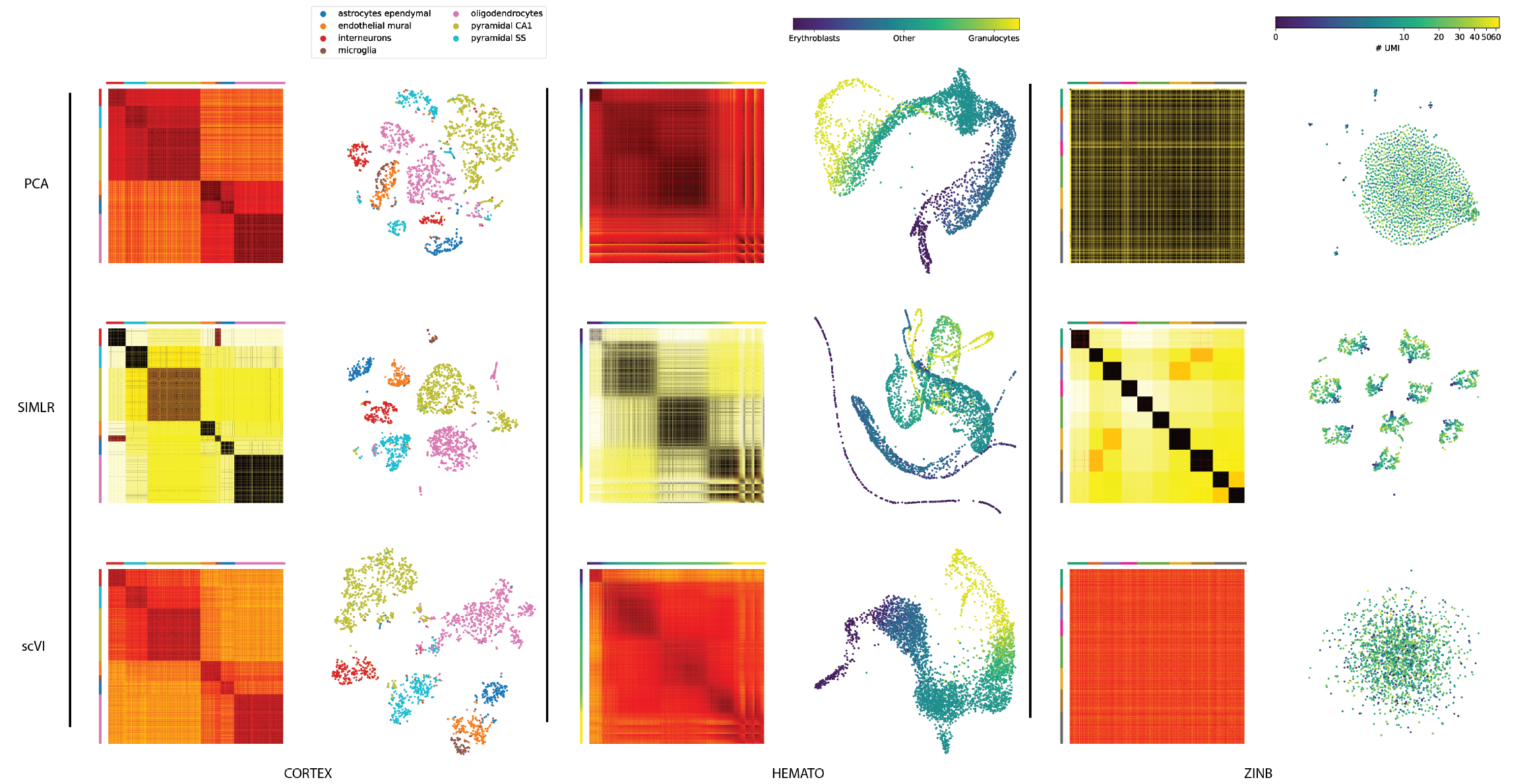
We apply scVI, PCA and SIMLR to three datasets (from right to left: CORTEX, HEMATO and a simulated ‘’noise” dataset sampled iid from a fixed zero inflated negative binomial (ZINB) distribution). For each dataset, we show a distance matrix in the latent space as well as a two-dimensional embedding of the cells. Distance matrices: the scales are in relative units from low to high similarity (over the range of values in the entire matrix). For CORTEX and HEMATO, cells in the matrices are grouped by their pre-annotated labels, provided by the original studies (for the CORTEX dataset, cell subsets were ordered by the hierarchical clustering in the original study). For ZINB, the color in the distance matrices is determined by the clusters called by SIMLR on this data. Embedding plots: each point represents a cell and the layout is determined either by tSNE (CORTEX, ZINB) or by a 5-nearest neighbors graph visualized using a Fruchterman-Reingold force-directed algorithm (HEMATO; see Supplementary Figure 15d for the original embedding for SIMLR). For CORTEX and HEMATO, the color scheme in the embeddings is the same as in the distance matrices. For ZINB, the colors reflect the number of UMI in each cell (see Supplementary Figure 15a-c for coloring of cells according to SIMLR clusters)

Finally, there may be the case of lack of structure, where the data is almost entirely dominated by noise. To explore this setting, we generated a noise dataset, sampled at random from a vector of zero-inflated negative binomial distributions. SIMLR erroneously reports eleven distinct clusters in this data, which are not perceived by any other method (Figure 3, Supplemental Figure 15abc).

Altogether, these results suggest that the latent space of scVI is flexible and describes the data well either as discrete clusters, as a continuum between cell state, or as structureless noise. scVI is therefore better suited than SIMLR in scenarios where the data does not necessarily fit with a simple structure of discrete subpopulations.

### 2.6 Controlling for batch effects

scVI explicitly accounts for the contribution of discrete nuisance factors of variation such as batch annotations in its graphical model. It does so by enforcing conditional independence between them and the inferred parameters.

Our model therefore learns gene expression bias that come from the batch effects and provides a parametric distribution that is disentangled from these technical effects, thus ideally reflecting the relevant biological variation. We evaluate the performance of scVI in correcting for batch effects using the RETINA dataset, which consists of two batches. We measured the entropy of batch mixing across the *K*-nearest neighbors graph (Methods 4.7) with the ideal expectation of a uniform representation of batches (i.e., maximum entropy) in any local neighborhood. We also measure the average silhouette width; with no batch bias, batches should overlap perfectly and exhibit a null silhouette width. Our results (Figure 4 and Supplementary Figure 8d) demonstrate that in this dataset scVI aligns the batches well, while maintaining a tight representation of pre-annotated subpopulations. Its performance in this regard is better than that of SIMLR as well as a more standard pipeline of batch correction: ComBat [34] followed by Principal Component Analysis (Methods 4.6). Notably, we performed a similar analysis with the PBMC dataset, which consists of cells from two donors. However, this data seemed to have very little batch effect to begin with (our metrics are averaged across all cell-types) and thus less informative for the purpose of this evaluation (Supplementary Figure 8a).

**Figure 4:**
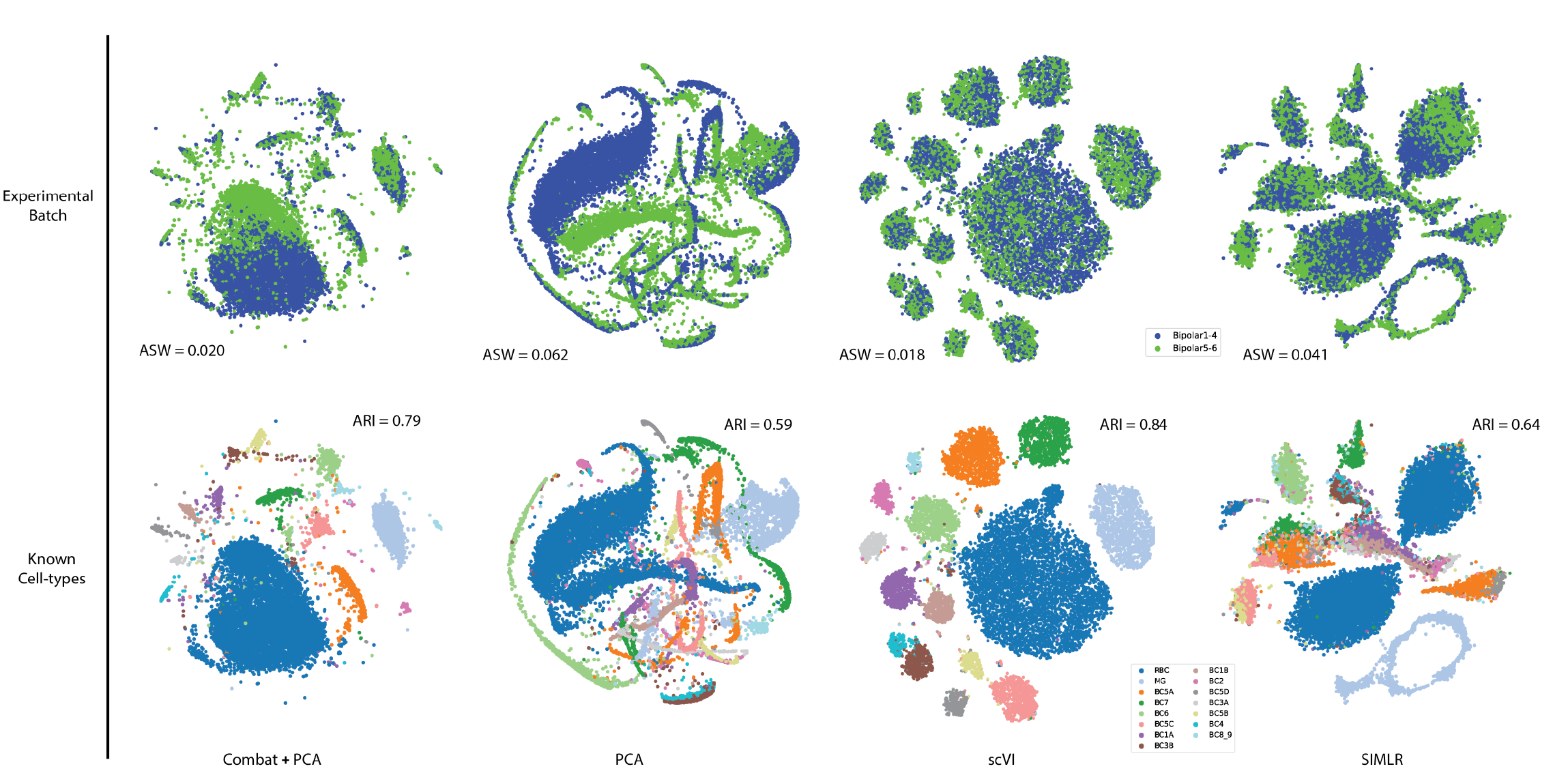
Batch effect removal with scVI on the RETINA dataset. We visualize the trade-off between batch removal and clustering. Embedding plots for the first three methods (PCA with COMBAT, PCA, scVI) were generated by applying tSNE on the respective latent space. For SIMLR, we used the tSNE coordinates provided by the program. In the upper part of the figure, the cells are colored by batch. In the lower part, cells are colored by the annotation of subpopulations, provided in the original study [30]. For each algorithm, we report the average silhouette width for the batches (ASW; lower is better) and the adjusted rand index for the clustering (ARI; higher is better). For SIMLR, the number of clusters was set to the number of pre-annotated subpopulations (*n* = 15). For PCA, we used the top 100 principal components (the top 10 resulted in no discernible structure).

### 2.7 Differential expression

Identifying genes that are differentially expressed between two subpopulations of cells is an important application of our generative model. The Bayesian model in scVI makes hypothesis testing straightforward. For clarity, through-out this section we assume that the cells are sequenced in the same batch *s*. (Methods 4.3 describes the general case.) For each gene *g* and a pair of cells (*z_a_*, *z_b_*) with observed gene expression (*x_a_, x_b_*), we consider two mutually exclusive hypotheses: 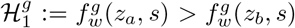 or 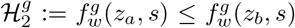. Evaluating which hypothesis is more probable amounts to evaluating a Bayes factor [35]:

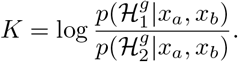

A Bayes factor is a Bayesian generalization of the p-value. Its sign indicates which of 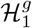 and 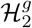 is more likely. Its magnitude is a significance level and throughout the paper, we consider a Bayes Factor as strong evidence in favor of a hypothesis if |*K| >* 3 [36] (equivalent to an odds ratio of *exp*(3) ≈ 20).

The posterior probability for each hypothesis can be approximated by integrating against the variational distribution:

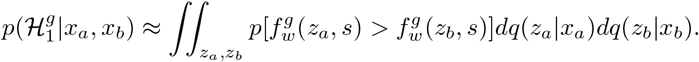

Sampling *z_a_* and *z_b_* from our variational distribution lets us approximate the integral with arbitrary precision. Since we model the cells as i.i.d., we can average the Bayes factors across randomly sampled cell pairs, one from each subpopulation. The average Bayes factor is a low-variance estimate of whether cells from one subpopulation tend to express *g* at a higher frequency.

We demonstrate the robustness of our method by repeating the entire evaluation process and comparing the results (Figure 5ab). We also ensure that our Bayes factor are well calibrated by running the differential expression analysis across cells from the same cluster and making sure no genes reach the signifiscance threshold (Supplementary Figure 12e).

**Figure 5:**
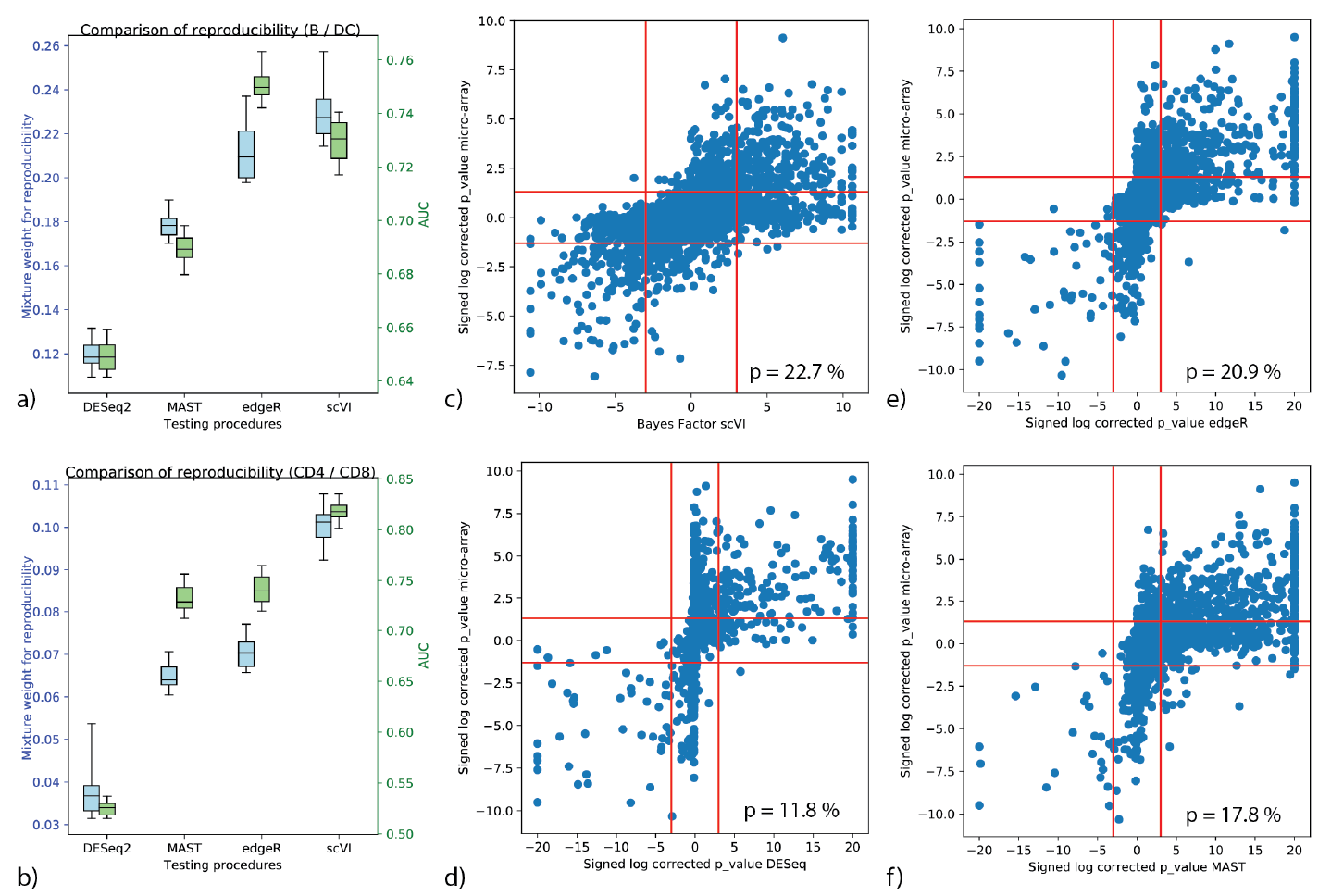
Benchmark of differential expression analysis using the PBMC dataset, based on consistency with published bulk data. (a, b) Evaluation of consistency with the irreproducible discovery rate (IDR) [37] framework (blue) and using AUROC (green) is shown for comparisons of B cells vs Dendritic cells (a) and CD4 vs CD8 T cells (b). Error bars are obtained by multiple subsampling of the data to show robustness. (c) through (f): correlation of significance levels of differential expression of B cells vs Dendritic cells, comparing bulk data and single cell. Points are individual genes. Bayes factors or p-values on scRNA-seq data are presented on the x-axis; Microarray-based p-values are depicted on the y-axis. Horizontal bars denotes significance threshold of 0.05 for corrected p-values. Vertical bars denotes significance threshold for the Bayes factor of scVI (c) or 0.05 for corrected p-values for DESeq2 (d), edgeR (e), and MAST (f). We also report the median mixture weight for reproducibility *p*. (Higher the better.)

To evaluate scVI as a tool for differential expression, we used the PBMC dataset along with its classification of cells into well studied subtypes of hematopoietic cells, for which reference bulk expression data is available.

We compare scVI to three widely used methods: DESeq2 [17], MAST [19] and edgeR [18]. To facilitate the evaluation, we defined a reference set of differentially expressed genes using publicly available bulk expression datasets. Specifically, we assembled a set of genes (Methods 4.7) that are differentially expressed between human B cells and dendritic cells (microarrays, n=10 in each group [38]) and between CD4+ and CD8+ T cells (microarrays, n=12 in each group [39]). We apply all four methods in these two differential expression tasks (using the respective clusters of cells) and evaluate the consistency with the reference data using two scores. For the first score, we assign each gene with a label of DE or non-DE based on their p-values from the reference data (genes with a corrected p-values under 0.05 to be positive and the rest negatives) and then these labels to compute AUROC for scVI and each of the benchmark methods.

Since defining the labels requires a somewhat arbitrary threshold, we use a second score that evaluates the reproducibility of gene ranking (bulk reference vs. single cell), considering all genes, using the irreproducible discovery rate (IDR) [37]. Considering the AUROC metric, scVI is the best performing method in the T cell comparison, while edgeR outperforms scVI by a smaller margin in the B vs. Dendritic cell comparison. Focusing on the proportion of genes with reproducible rank as fitted by IDR, scVI is the best performing method (Figure 5, Supplementary Figure 12a-d).

### 2.8 Capturing technical variability

To further interpret the fitted models, we study the extent to which they capture technical variability. We focus on datasets that were generated by 10x, as they share the same set of cell quality metrics (generated by cell ranger; see Methods 4.5) and can thus provide reproducible insights about the relationship between our parameters and library quality. Additionally, we required our test datasets to have pre-annotated subpopulations, with the assumption that each subpopulation consists primarily of cells of the same type, thus decreasing the extent of biological heterogeneity (e.g., in total mRNA content). The two datasets that fit these requirements, which we report next are the PBMC and BRAIN-SMALL.

In each case, we trained the model on the entire dataset and then investigated each pre-annotated subpopulation separately. As a general rule, we find that variation in library size correlates strongly with the cell-specific scaling factor, which is to be expected by the definition of the model. Figure 6a depicts these results for the CD14+ monocytes subpopulation in the PBMC data (notably, the negative curvature on the plot can be explained by shrinking the values towards the mean due to the use of prior). Another important nuisance factor to be considered is the limitation in sensitivity, which results in exacerbated amount of zero entries. In principle, zero entries can be captured by two different components of our model: the negative binomial and the “inflation” of zeros added to it with a Bernoulli distribution. Evidently, the expected number of zeros generated by the negative binomial for each cell correlates strongly with the library size and its proxies (e.g., the number of detected genes or the number of reads per UMI; Figure 6cd, Supplementary Figure 13d). This result can be explained by our definition of the negative binomial mean, which is the predicted frequency of expression 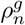 scaled by the respective library size *ℓ_n_*.

**Figure 6:**
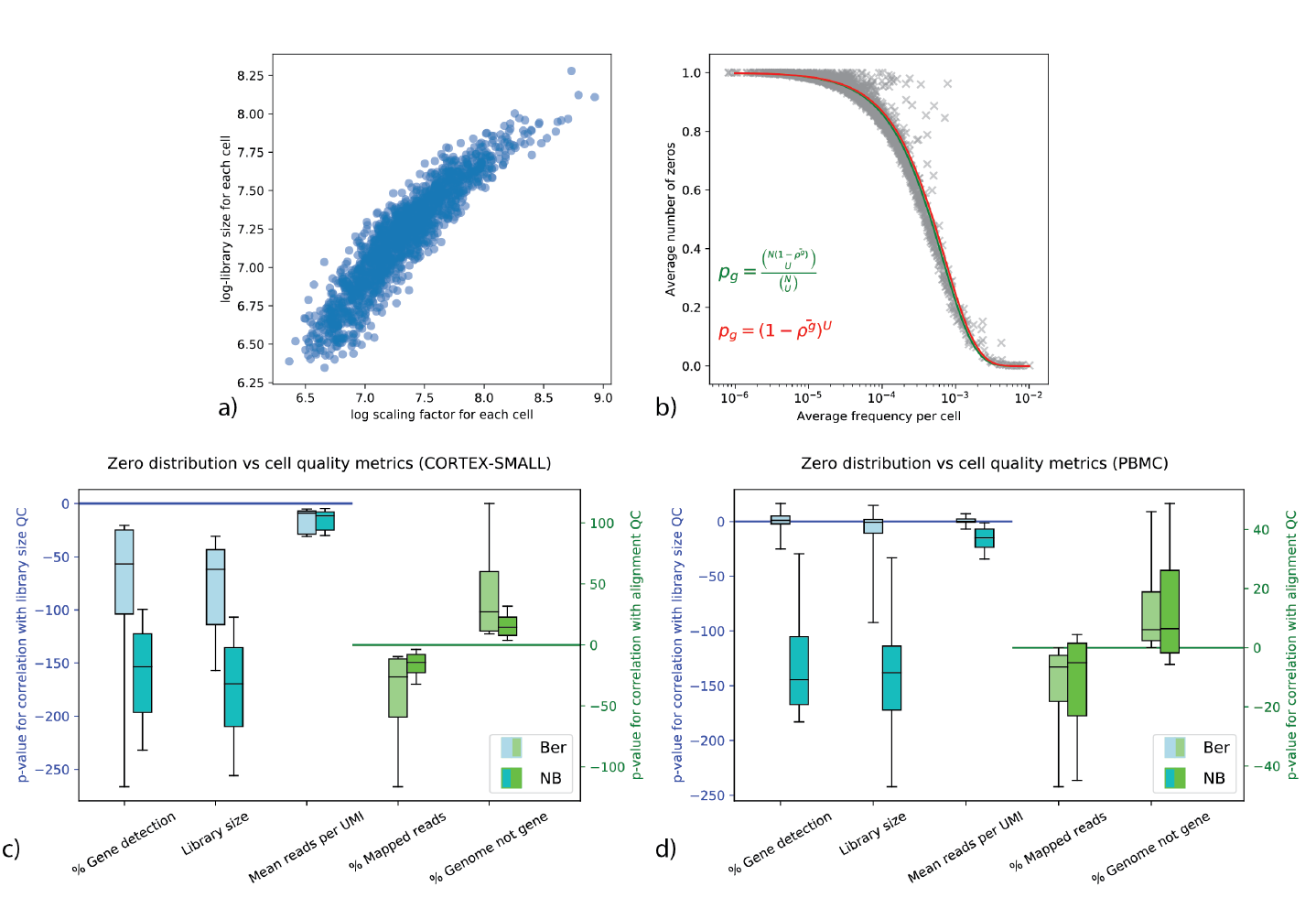
Capturing technical variability with the parameters of scVI. Data for panels a-b is based on the CD14+ cell subpopulation in the PBMC dataset (a) Scatter plot for each cell of inferred scaling factor by scVI against library size. (b) The frequency of observed zero values versus the expected expression level, as produced by scVI. Each point represents a gene *g*, where the x-axis is 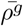 - the average expected frequency per cell (for gene *g*, average over 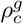 for all cells *c* in the subpopulation), and the y-axis the is observed percentage of cells that detect the fee (UMI>0). The cyan curve depicts the probability for selecting zero transcripts from every gene as a function of its frequency, assuming a simple model of sampling *U* molecules from a cell with *N* molecules at random without replacement, where *U* = 1398 is the average number of UMIs in the subpopulation. The hypergeometric distribution (inset) requires the average number of transcripts per cell (*N*). Notably the curve converges for values larger than 20*k* (here, we set *N* = 10*k*). Indeed, when *N → ∞* the process converges to a binomial selection procedure, which is depicted by the red line. (c, d) Signed log p-values for testing correlations between the zero probabilities from the two distributions (negative binomial, Bernoulli) and quality control metrics across five random initializations of scVI and all subpopulations of the PBMC and the BRAIN-SMALL datasets.

**Figure 7:**
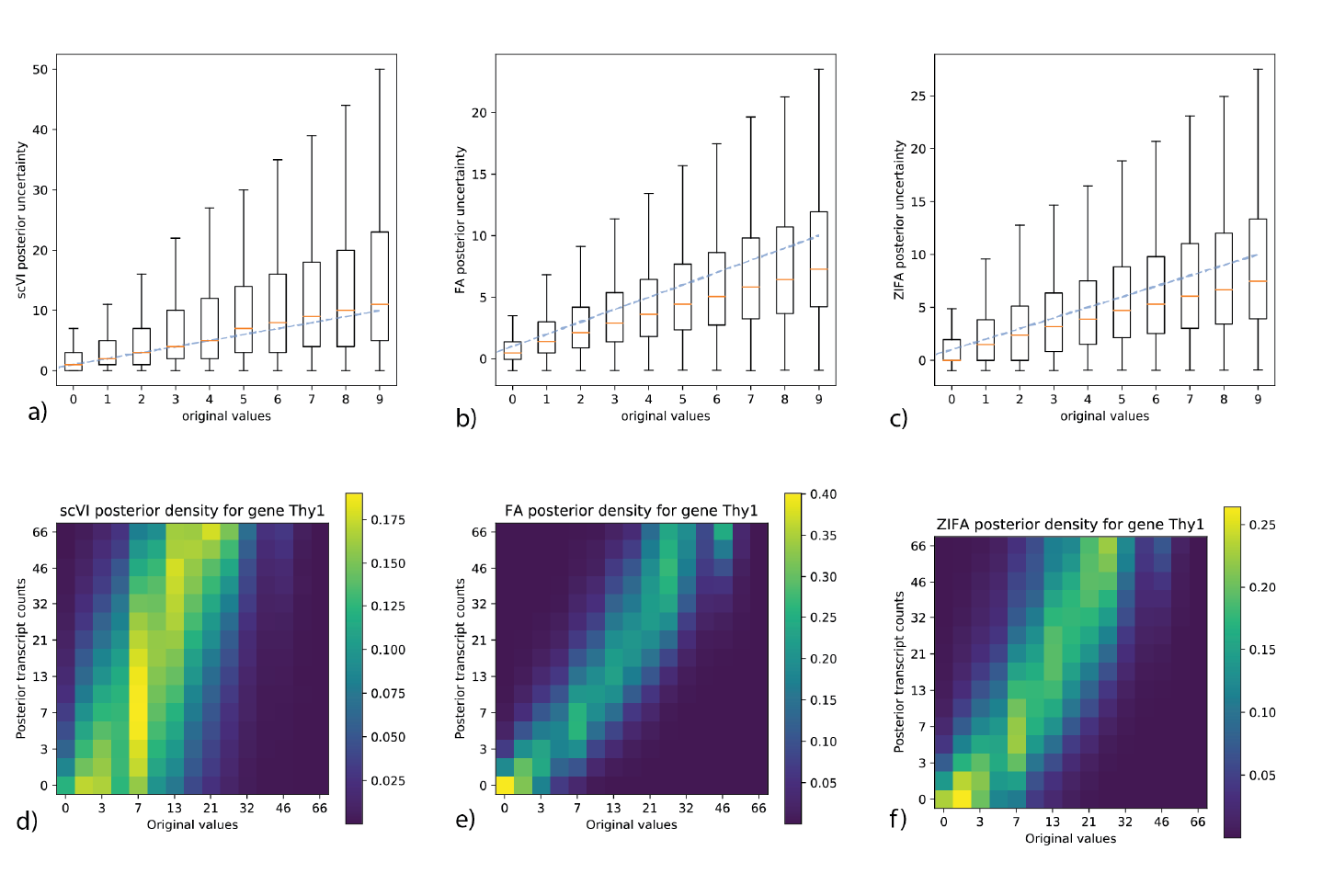
Posterior analysis of generative models on the CORTEX dataset. Panels (a-c) depict the observed counts of randomly selected entries of the data matrix (X axis) and their posterior uncertainty (Y axis) by sampling from the variational posterior (scVI) or the exact posterior (FA, ZIFA). Panels (d-f) represents the observed counts of a representative gene, Thy1, in the CORTEX dataset. Data is presented across all cells (X axis) against the posterior expected counts produced by scVI, ZIFA and FA respectively (Y axis). The values in each axis have been divided into 20 bins and the color scale reflects the proportion of cells in each pair of bins. By definition, the uncertainty of FA is independent of the input value and tight around the observed count. ZIFA can generate zero and puts realistically more weight in this area. scVI’s posterior is more complex, able to generate zero for low UMI values but also able to generate high UMI values when the original count observed was only of intermediate intensity.

**Figure 8:**
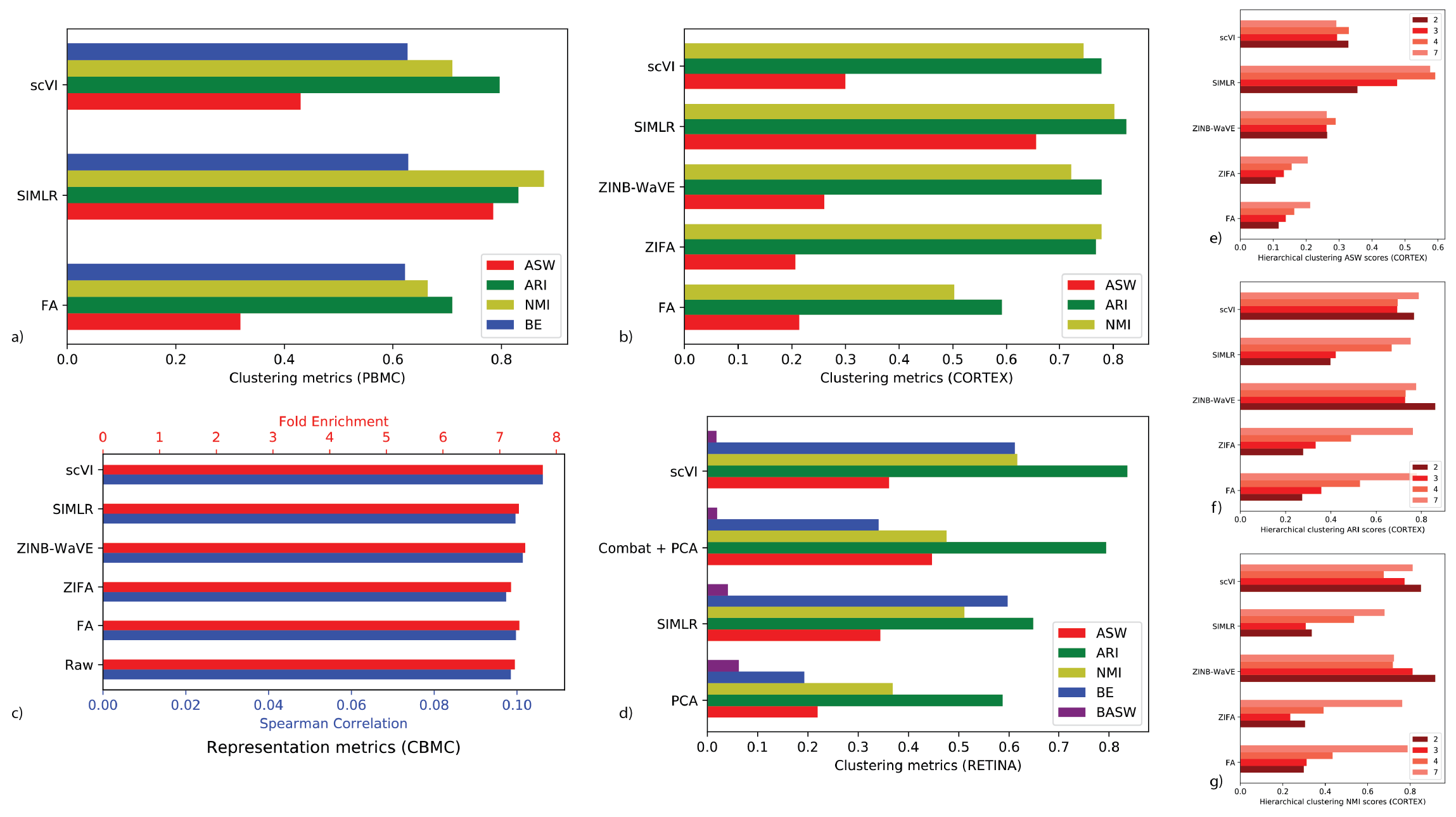
We investigate how scVI latent space can be used to cluster the data and report benchmarking across datasets for state-of-the-art methods. The first four panels depict the results for the (a) PBMC dataset (b) CORTEX (c) CBMC and (d) RETINA. ASW: average silhouette width of pre-annotated subpopulations (higher is better), ARI: adjusted rand index (higher is better), NMI: normalized mutual information (higher is better), BE: batch mixing entropy (higher is better), BASW: average silhouette width of batches (lower is better). Panels (e-g) depict the performance of clustering metrics for different depth of the hierarchical clustering in the CORTEX data, computed in the original publication [28]. The numbers in the legend indicate the number of clusters at the given depth.

**Figure 9:**
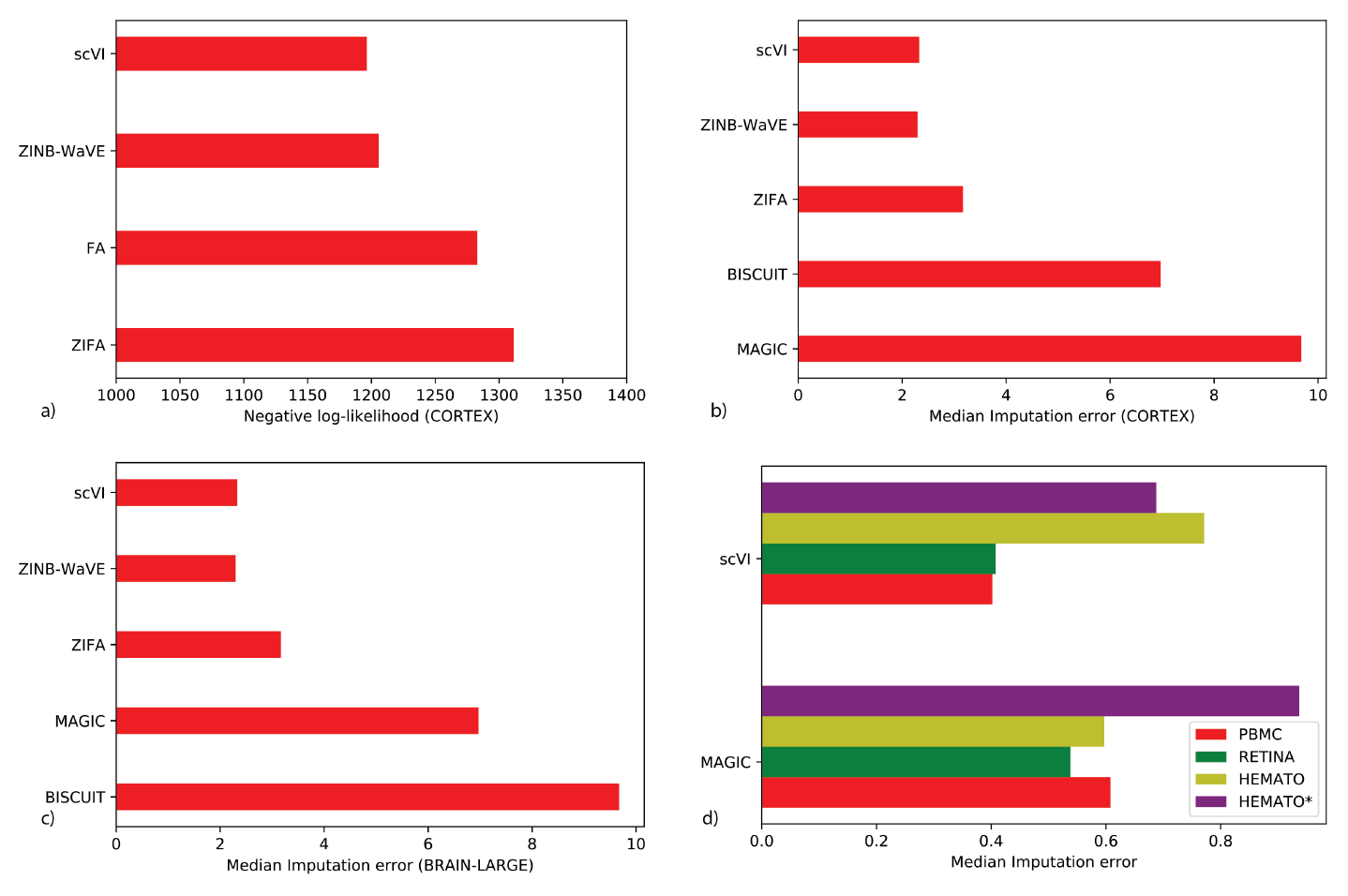
(a) Log-likelihood results on the CORTEX dataset. (b) through (d): we investigate how scVI latent space can be used to impute the data (with the uniform perturbation scheme) and report benchmarking across datasets for state-of-the-art methods.

**Figure 10:**
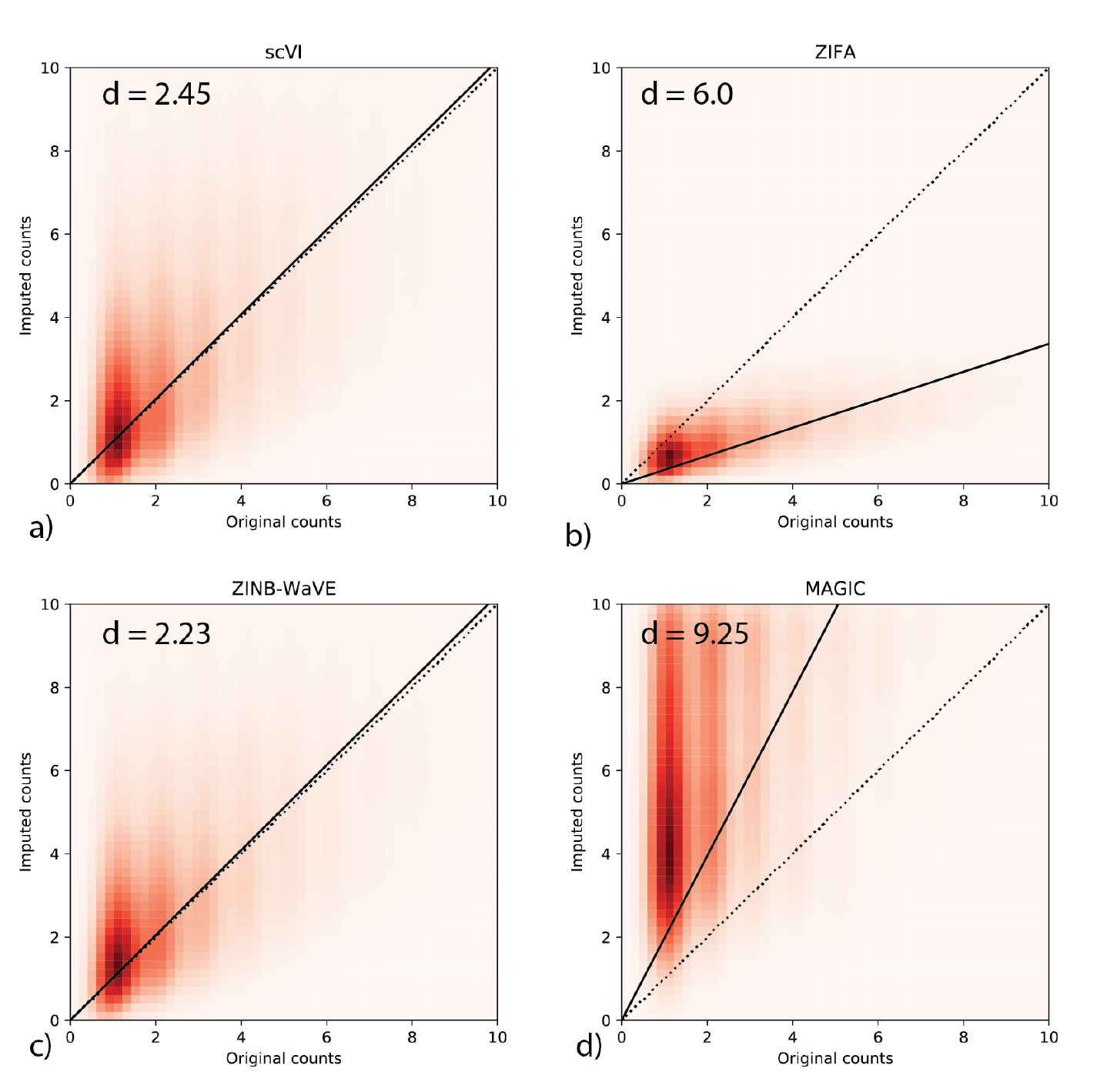
Imputation of scVI on the CORTEX dataset. Models are trained on a binomial-down-sampling corrupted dataset (see Methods). The heatmaps denotes density plots of imputed values scVI, ZIFA, MAGIC and ZINB-WaVE on a down-sampled version vs original values prior to down-sampling. The reported score *d* is the median imputation error across all the hidden entries (Lower is better; see Methods).

**Figure 11:**
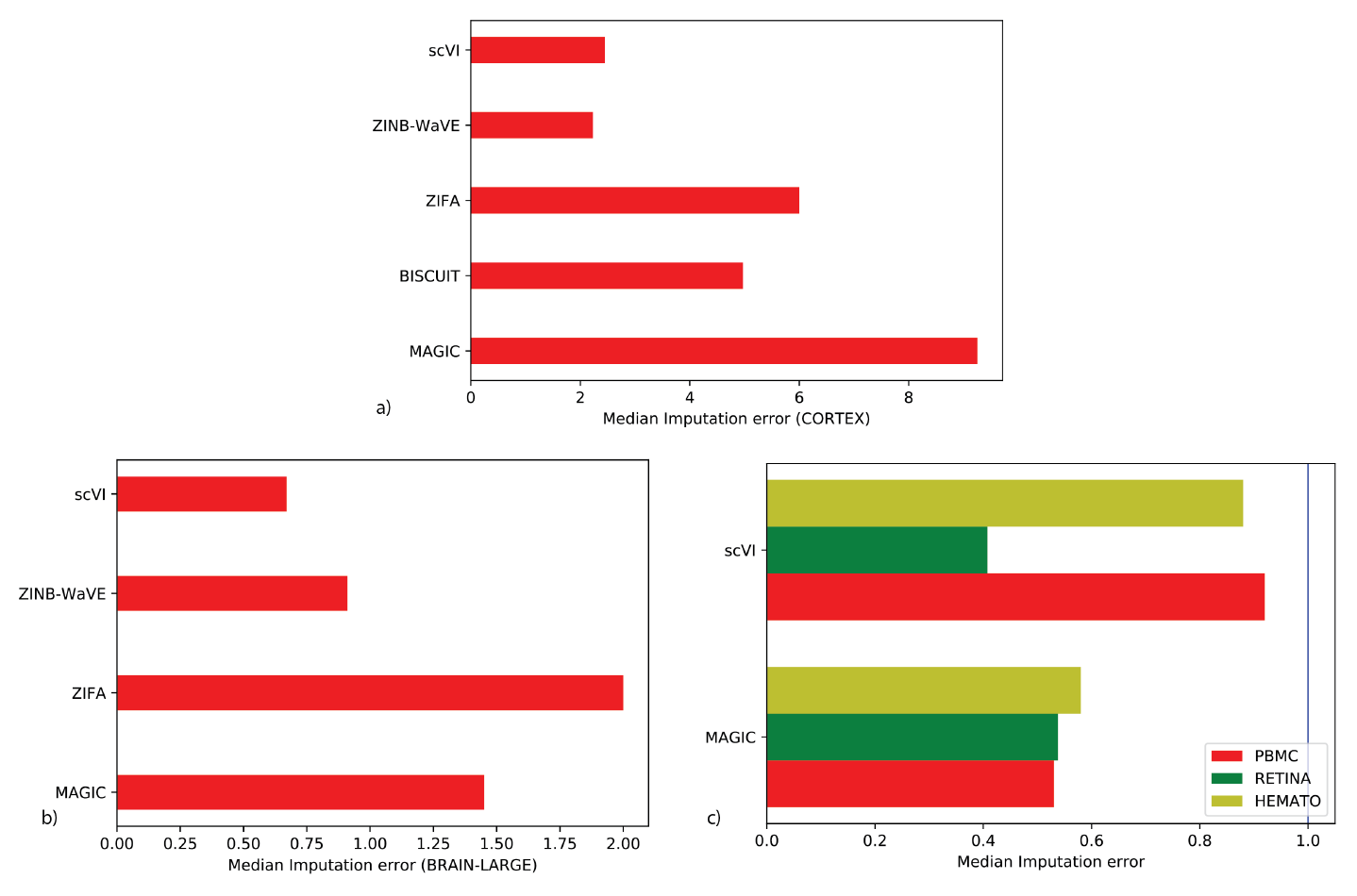
(a) through (c): we investigate how scVI latent space can be used to impute the data (with the binomial perturbation scheme) and report bench-marking across datasets for state-of-the-art methods.

**Figure 12:**
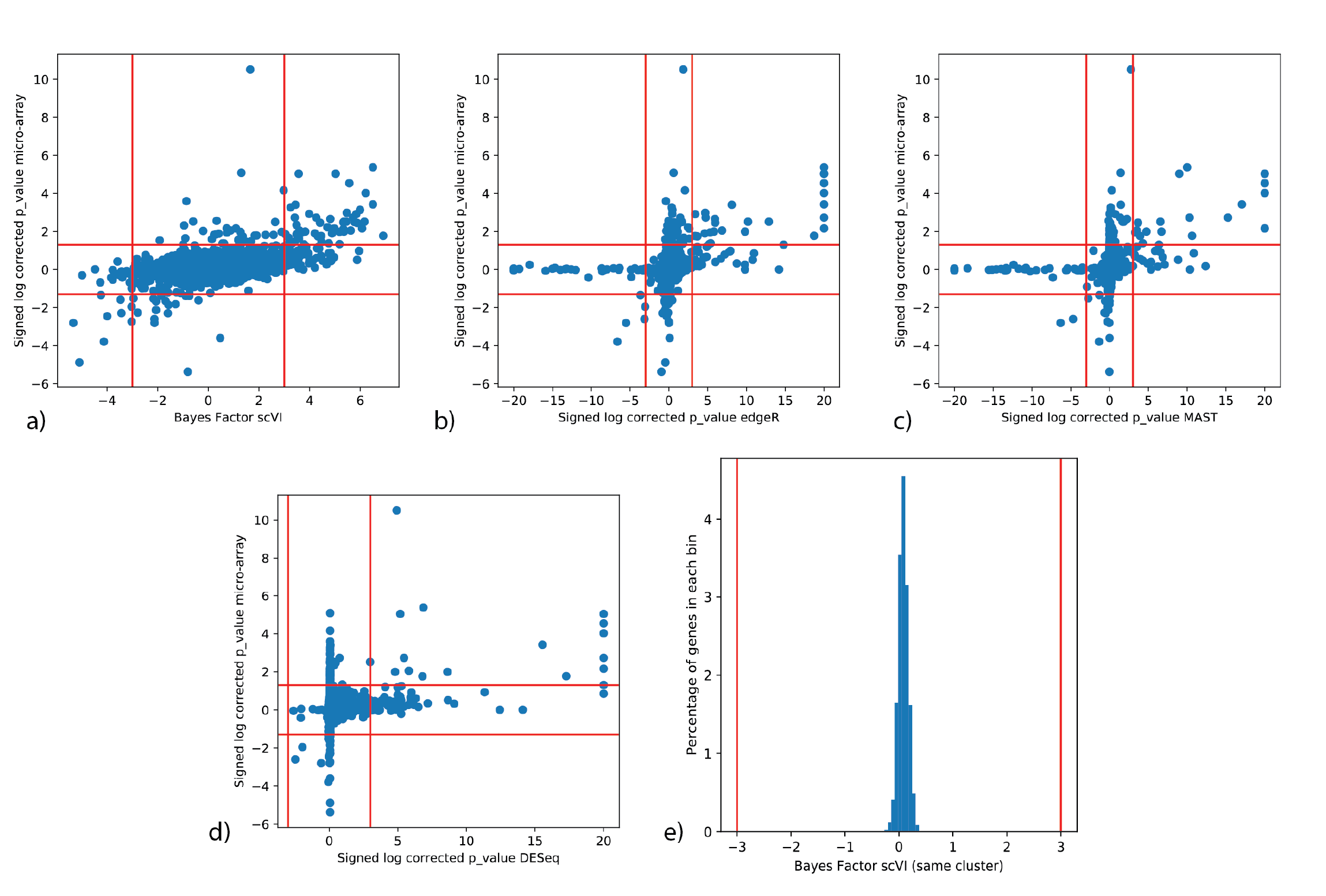
Differential expression with scVI on the PBMC dataset. (a) (b) (c) (d) p-values of microarray against p-values or Bayes factor for CD4 / CD8 comparison. In that order, scVI, edgeR, MAST, DESeq2 (g) Bayes factor of scVI when applying DE to two sets of random cells from the same cluster.

**Figure 13:**
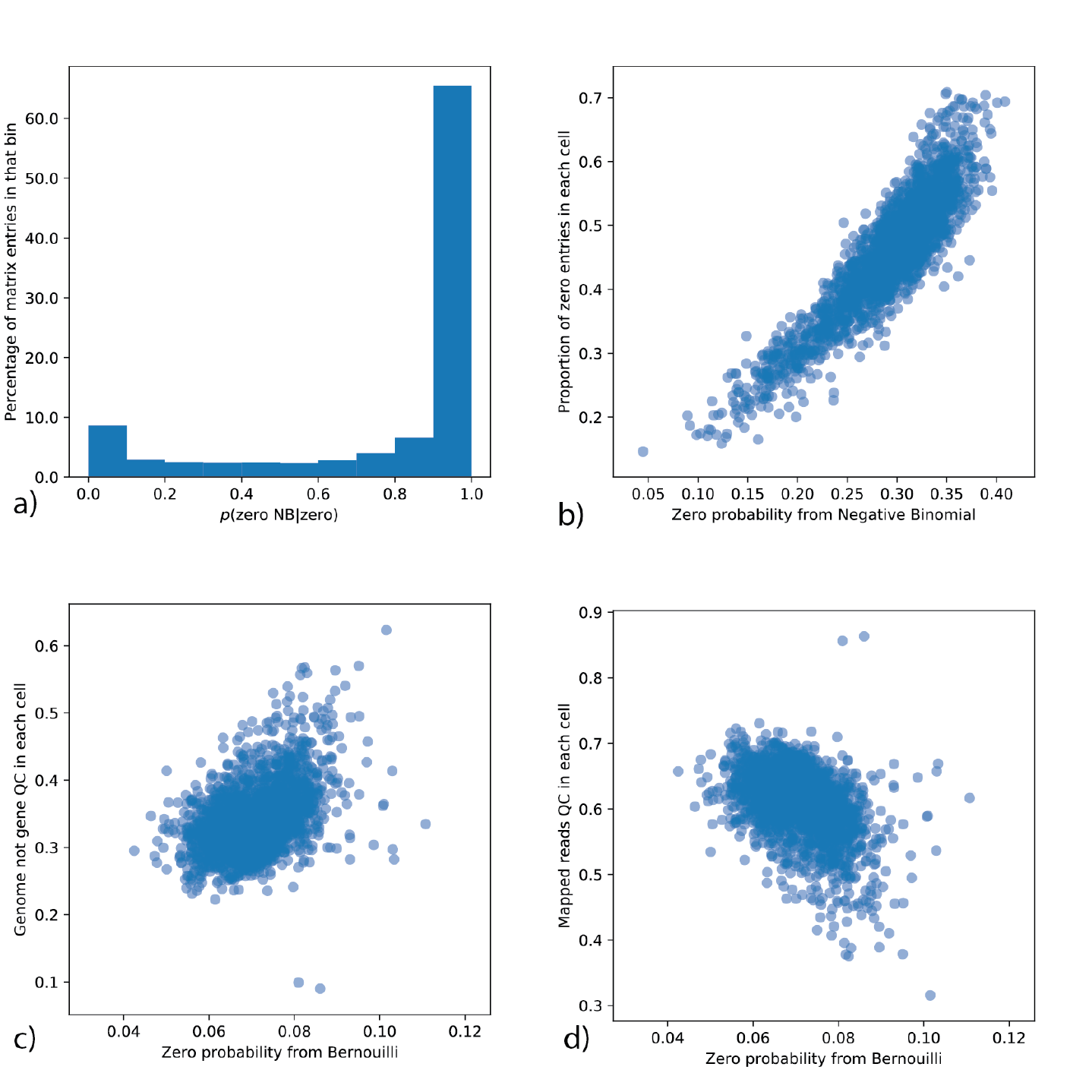
The generative distributions of scVI. This study focuses on a particular subpopulation of the BRAIN-SMALL dataset (a) To sustain that most of the zeros in the data comes from the Negative Binomial, we plot for each entry of the count matrix (percentage in Y axis) the probability that a given zero comes from the NB conditioned on having a zero (X axis). (b) Number of gene detected vs negative binomial zero probability averaged across all genes. (c) Genome_not_gene vs Bernoulli zero probability averaged across all genes. (d) Mapped_reads vs Bernoulli zero probability averaged across all genes.

**Figure 14:**
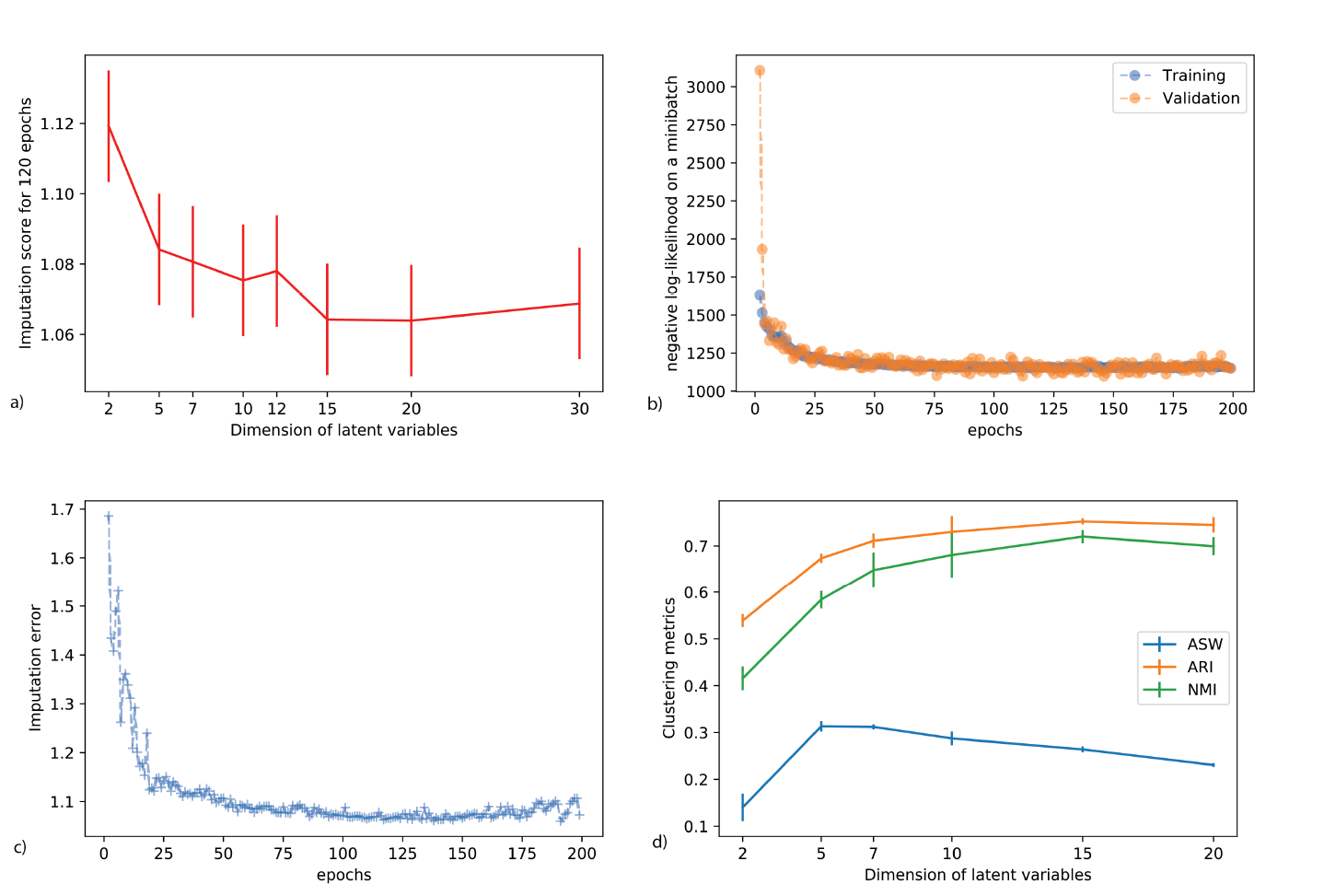
Robustness analysis for scVI. (a) Imputation score on the BRAIN-LARGE dataset across multiple random initialization, training and dimension of the latent space. (b) Visualization of scVI numerical objective function during training on the BRAIN-LARGE dataset. This shows our model do not over fit and has a stable training procedure. (c) Imputation score as a function of the number of epochs on the BRAIN-LARGE dataset. This figure also show stability across posterior sampling since there is not much change in the parameters between two subsequent epochs. (d) Clustering metrics on the CORTEX dataset across multiple initializations and dimensions for the latent space.

**Figure 15:**
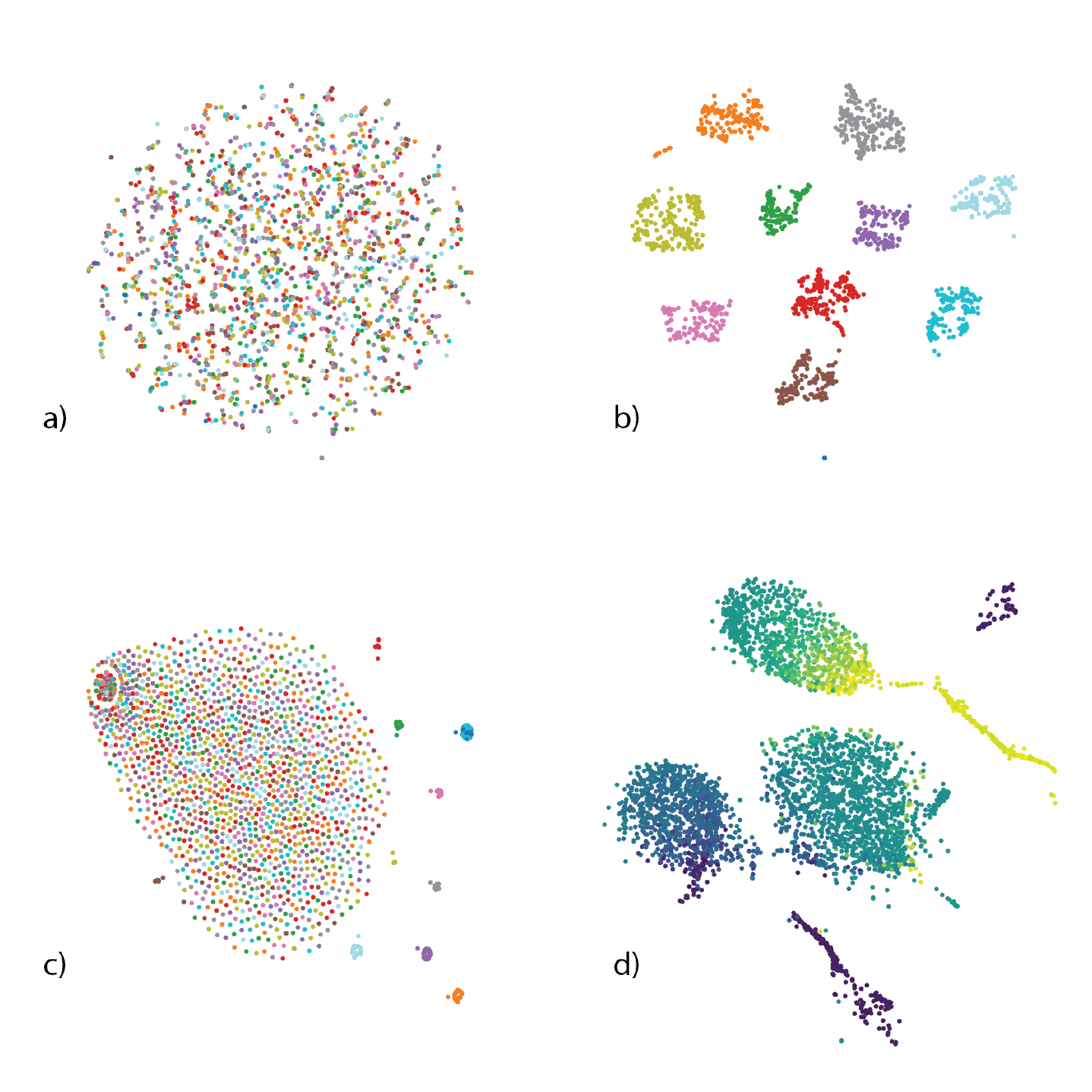
Details for the clustering panel. For the random data, we obtain the labels that order the cell-cell similarity matrices by a k means clustering on SIMLR latent space. (a) scVI latent space with SIMLR labels. There is no structure. (b) SIMLR latent space with SIMLR labels. (c) PCA latent space with SIMLR labels. (d) SIMLR tSNE on the HEMATO dataset. We prefer to visualize the SIMLR embedding on a kNN graph since even tSNE would loose the continuum structure of the dataset.

**Figure 16:**
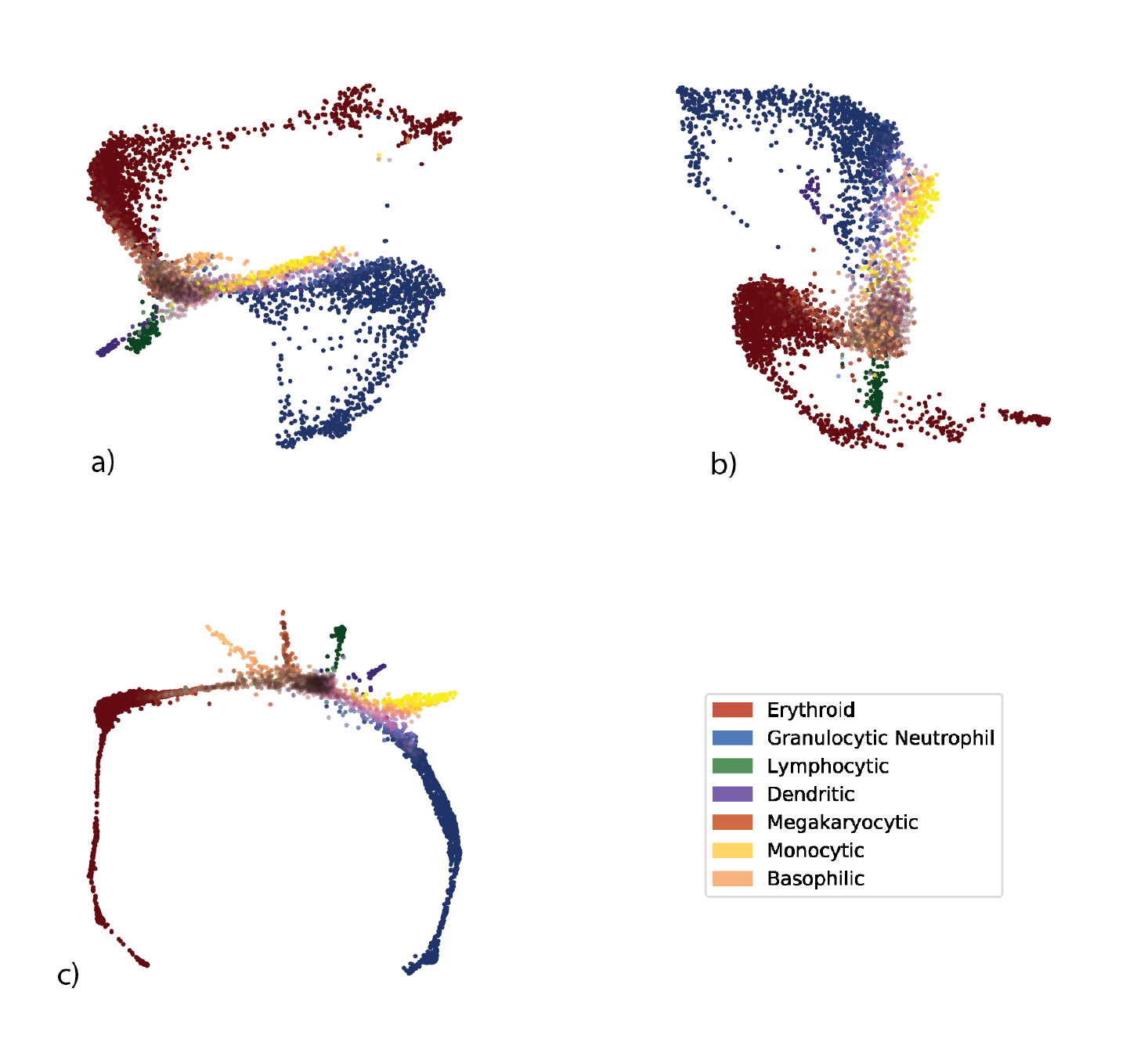
Additional comparison of scVI and PCA on the HEMATO dataset. All scatter plots picture the embedding of a 5-nearest neighbors graph of a latent space. Cells positions are computed using a force-directed layout (a) denotes a reduction to 60 pcs as in the original paper. (b) denotes the output of a scVI in dimension 60. As the dimension is sensibly different from other experiments, the warm-up schedule (which governs how the prior on *z* is enforced) was adjusted. (c) denotes the figure from the main paper. To recover all the differentiation paths, the authors performed several operations on the *K*-nearest neighbors graph that we did not reproduce in this analysis. We instead visualize the graph before the smoothing procedure.

The remaining question is therefore: what is the relationship between the expected frequency of expression 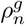 and the observed zeros? A simple model would be a random process of sampling genes from each cell in manner proportional to their frequency, and with no added bias (e.g., in capture efficiency). To explore this, we plot for every gene the mean expected frequency against the percent of detecting cells in a subpopulation of interest (Figure 6b). The resulting trend supports this simple model as it closely fits with the zero probability of a hypergeometric distribution—namely random sampling of molecules without replacement.

Interestingly, the number of additional zeros induced by the Bernoulli random variable in each cell is less correlated with library size, and instead correlates with metrics of alignment rate (Figure 6cd, Supplementary Figure 13cd). These metrics are not necessarily coupled to size, but may reflect other technical factors such as contamination or the presence of degraded mRNA. However, we observed that most zero values in the data can be explained by the negative binomial component alone (Supplementary Figure 13a). Taken together, these results therefore corroborates the idea that most zeros, at least in the datasets explored here, can be explained by low (or zero) “biological” abundance of the respective transcript, exacerbated by limited sampling.

## 3 Discussion

Our study focuses on an important need in the field of single cell RNA-seq - namely accounting for confounding factors and measurement uncertainty in tertiary analysis tasks (e.g., clustering, differential expression, annotation) through a common, scalable statistical model. To achieve this, we developed scVI - a hierarchical Bayesian model, which makes use of neural networks to provide a complete probabilistic representation of single cell transcriptomes. We demonstrated that scVI provides a computationally efficient and “all inclusive” approach to denoising and analyzing gene expression data, and showcase its performance in downstream tasks including dimensionality reduction, imputation, visualization, batch-effect removal, clustering and differential expression.

The scVI procedure takes as input a matrix of counts and therefore does not need a preliminary step of normalization. Instead, it learns a cell-specific scaling factor as a hidden variable of the model, with the objective of maximizing the likelihood of the data (as in [10, 9, 40]), which is more justifiable than correcting *a posteriori* the observed counts [5]. Furthermore, scVI explicitly accounts for the contribution of discrete nuisance factors, such as batch annotations, by enforcing conditional independence between them and the (inferred) parameters that govern gene expression distributions. Since this correction step is performed via the mild modeling assumption of conditional independence, scVI can reasonably integrate and harmonize multiple datasets. Further modeling would be needed for more intensive usage of batch removal (number of batches/ datasets ≥ 20) and is left as future research direction.

An important distinguishing feature of scVI is its scalability. Unlike the benchmark methods, scVI is capable of efficiently processing very large datasets, with up to a million cells explored in this study[21, 20]. To achieve this high level of scalability while ensuring good fit to the data, we designed an efficient procedure to learn the parameters of our graphical model. Importantly, exact Bayesian inference is in most cases not tractable for these kinds of models. Furthermore, until recently, even variational inference was rarely applied to such models without restrictive “conditional conjugacy” properties. To address this, we use a stochastic optimization procedure that samples our approximation of the posterior distribution (as well as “minibatches” of our dataset), allowing us to efficiently perform inference with arbitrary models, including those with conditional distributions specified by neural networks [27].

Notably, since our procedure has a random component and since it optimizes a nonconvex objective function, it may give alternative results from different initializations. To address this concern, we demonstrate the stability of scVI in terms of its objective function, as well as imputation and clustering (Supplemental Figure 14). Another related issue is that if there are few observations (cells) for each gene, the prior (and the inductive bias of the neural network) may keep us from fitting the data closely. Indeed, in the case of datasets such as HEMATO [31] where the number of cells is smaller than the number of genes, some procedure to pre-filter the genes may be warranted. Another approach that can help make scVI applicable to smaller data sets (hundreds of cells) and which we intend to explore, is to utilize techniques such as Bayesian shrinkage [17] or regularization and second order optimization with larger mini-batch size / full dataset [9]. We do however show that for a range of datasets of varying sizes, scVI is able to fit the data well and capture relevant biological diversity between cells.

From a system perspective, single-cell RNA-seq analyses paradoxically benefit from the abundance of zero values, as it allows one to store the data in sparse (rather than full matrix) format. A sparse matrix with one million cells and ten thousand genes would represent around 7.5 GB assuming one percent sparsity. On the other hand, the output of batch-corrected data is not sparse and therefore potentially very large (for 1M cells, approximately 75 GB). While it performs batch correction, scVI still provides a compact representation of the complete data, as it requires only the latent space and the specification of the model (overall less than 1G memory footprint, assuming 10 latent variables). One obvious drawback of such compressed representation (apart from the potential loss of information) is that gene expression values need to be computed on the fly; however this can be done very efficiently in a single-pass through the generative networks (approximately 10 seconds for generating the *ρ* matrix for a test dataset of 500k cells and 8k genes, with the same hardware specification used throughout this paper). This property makes scVI a good baseline to be used by interactive visualization tools [41, 42, 43].

Looking forward, it is important to note that the model of scVI is very general and therefore provides a proper statistical framework for other forms of scRNA-seq analysis, not explored in this manuscript, such as lineage inference [2] or cell state annotation [1, 7]. Furthermore, as the scale and diversity of single-cell RNA-seq increase, we expect tools such as scVI to be in great demand, especially in cases where there is interest in harmonizing datasets in a manner that is scalable and conducive to various forms of downstream analysis [21]. Indeed, one subsequent research direction would be to merge multiple datasets from a given tissue to build a generative model with biological annotations of cell-types or phenotypical conditions in a semi-supervised fashion. That would allow researchers to “query” the generative model with a new dataset in an online fashion in order to “retrieve” previous biological information, which would allow verification of reproducibility across experiments, as well as transfer of cell state annotations between studies.

## 4 Online Methods

### 4.1 The scVI probabilistic model

First, we present in more detail the generative process for scVI:

**Require:** constant prior for cell-specific scaling *ℓ_μ_, ℓ_σ_*

**Require:** fitted gene-specific inverse dispersion parameter *θ*

**Require:** fitted neural networks *f_w_, f_h_*

**Require:** observed batch ID *s_n_*

~~~
1: **for** cell *n* **do**
2:    Choose a low-dimensional vector *z_n_ ~* 𝒩 (0*, I*) describing the cell
3:    Choose a batch *s_n_* ∈ {1*..B*} from which the cell is sampled
4:    Choose a cell-scaling factor 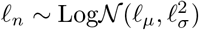
5:    **for** gene *g* in gene set *G* **do**
6:      Choose a normalized expression mean *w_ng_ ~* Gamma(*f_w_*(*z_n_*)*, θ*).
7:      Choose an expression level *y_ng_ ~* Poisson(*ℓ_n_w_ng_*).
8:      Choose a dropout event with *h_ng_ ~* Bernoulli(*f_h_*(*z_n_*))
9:      Apply dropout to expression level *y_ng_* and output *x_ng_*
10:   **end for**
11: **end for**
~~~

A standard multivariate normal prior for *z* is commonly used in variational autoencoders since it can be reparametrized in a differentiable way into any arbitrary multivariate Gaussian random variable [27], which turns out to be extremely convenient in the inference process (see Methods 4.2). *ℓ_μ_, ℓ_σ_* are set to be the empirical mean and variance of the log-library size per each batch. Let us note that the random variable *ℓ_n_* is not the log-library size (scaling the sampled observation) itself but a scaling factor who is expected to correlate strongly with log-library size (hence the choice for the parameters). Neural network *f_w_* is constrained during the inference to encode the mean proportion of transcripts expressed across all genes by using a softmax activation at the last layer. Neural network *f_h_* encodes whether a particular entry has been dropped out due to technical effects [11, 9].

All neural networks use dropout regularization and batch normalization. Each network has 1, 2, or 3 fully connected-layers, with 128 or 256 nodes each. The activation functions between two hidden layers are all ReLU. We use standard link function to parametrize the distributions parameters (exponential, logarithmic or softmax). Weights for some layers are shared between *f_w_* and *f_h_*. Throughout the paper, we use Adam as first order stochastic optimizer with ϵ = 0.01 fixed. A complete list of datasets, their properties, the applicable methods and a thorough list of hyperparameters is provided in Table 2. We optimize the objective function until convergence (usually between 120 and 250 epochs, where each epoch is a complete pass through the dataset. Let us note that bigger datasets require less epochs). The implementation for all of the analysis performed in this paper is available at https://github.com/romain-lopez/scVI-reproducibility. The reference implementation is available at https://github.com/YosefLab/scVI.

### 4.2 Fast inference via stochastic optimization

The posterior distribution combines the prior knowledge with information acquired from the data *X*. We cannot directly apply Bayes rule to determine the posterior because the denominator (the marginal distribution) *p*(*x_n_*|*s_n_*) is intractable. Making inference over the whole graphical model is not needed. We can integrate out the latent variables *w_ng_*, *h_ng_* and *y_ng_* by making sure the conditional *p*(*x_ng_|z_n_, ℓ_n_, s_n_*) has a closed-form density and is Zero-Inflated Negative Binomial (see Appendix A). Having simplified our model, we use variational inference [26] to approximate the posterior *p*(*z_n_, ℓ_n_|x_n_, s_n_*). Our variational distribution *q*(*z_n_, ℓ_n_|x_n_, s_n_*) is mean-field:

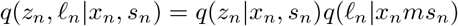

The variational distribution *q*(*z_n_|x_n_, s_n_*) is chosen to be Gaussian with a diagonal covariance matrix, mean and covariance are given by an encoder network applied to (*x_n_, s_n_*), as in [27]. The variational distribution *q*(*ℓ_n_|x_n_, s_n_*) is chosen to be log-Normal with scalar mean and variance also given by an encoder network applied to (*x_n_, s_n_*). The variational lower bound is

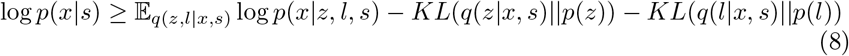

In this objective function, the dispersion parameters *θ_g_* for each gene are treated as global variables to optimize in a Variational Bayesian inference fashion. To optimize the lower bound, we use the analytic expression for *p*(*x|z, l, s*) and use analytic expressions for the Kullback-Leibler divergences. We use the reparametrization trick to compute low-variance Monte-Carlo estimates of the expectations’ gradients. Analytic closed-form for the Kullback-Leibler divergence and the reparametrization trick are only possible on certain distributions which multivariate Gaussians are a part of [27]. Now, our objective function is continuous and end-to-end differentiable, which allows us to use automatic differentiation operators. As indicated in [44], we use deterministic warmup and batch normalization during learning to learn an expressive model.

Our objective function is nonconvex and thus could give alternative results from different initializations. We show stability of our algorithm and its results in Supplemental Figure 14.

Since our model assumes cells are identically independently distributed, we can also benefit from stochastic optimization from sampling the training set. We then have an online optimization procedure that can handle massive datasets. At each iteration, we only focus on a small subset of the data randomly sampled (*M* = 128 data points) and need not go through the entire dataset. There is then no need to store the entire dataset in memory. Because the number of gene is limited in practice to a few tens of thousands, these mini-batches of cells fit easily into a GPU.

Since the encoder network *q*(*z|x, s*) might still produce output correlated with the bath *s*, we use a Maximum Mean Discrepancy (MMD) based penalty as in [45] to correct the variational distribution. For this paper, we however did not explicitly enforce the MMD penalty and just retained the conditional independence property that has shown to be efficient enough. It might be useful on other datasets.

### 4.3 Bayesian Differential Expression

For each gene *g* and a pair of cells (*z_a_*, *z_b_*) with observed gene expression (*x_a_, x_b_*) and batch ID (*s_a_, s_b_*), we can formulate two models of the world under which one of the following hypotheses is true

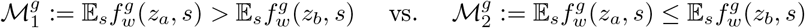

where the expectation 𝔼_*s*_ is taken with the empirical frequencies. Notably, we propose a hypothesis testing that do not to calibrate the data to one batch but will find genes that are consistently differentially expressed. Again, evaluating the likelihood ratio test for whether our datapoints (*x_a_, x_b_*) are more probable under the first hypothesis is equivalent to writing a Bayes factor:

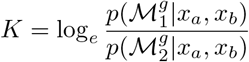

where the posterior of these models can be approximated via the variational distribution:

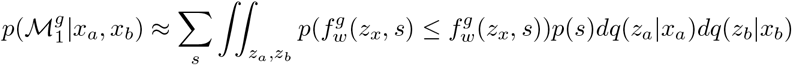

where *p*(*s*) designated the relative abundance of cells in batch *s* and all the measures are low-dimensional so we can use naive monte-carlo to compute these integrals. We can then use a Bayes factor for the test.

Since we assume that the cells are i.i.d., we can average the Bayes factors across a large set of randomly sampled cell pairs, one from each subpopulation. The average factor will provide an estimate to whether cells from one subpopulation tend to express *g* at a higher frequency.

### 4.4 Software implementation

We discuss numerical stability and parametrization of the ZINB distribution in Appendix B. Our model is implemented in Python and TensorFlow. A functional code can be found at https://github.com/YosefLab/scVI

### 4.5 Datasets and preprocessing

Below we describe all the datasets and the preprocessing steps used in the paper. A star after the dataset name means we used it as an auxiliary dataset and do not use it for general benchmarking but in order to make a specific point in the paper. The only case where we subsample the data multiple times is for the BRAIN-LARGE dataset. However, we just use one instance of it to report all possible scores (further details in Table 2).

#### CORTEX

The Mouse Cortex Cells dataset from [28] contains 3005 mouse cortex cells and gold-standard labels for seven distinct cell types. Each cell type corresponds to a cluster to recover. We retain top 558 genes ordered by variance as in [10].

#### PBMC

We considered scRNA-seq data from two batches of peripheral blood mononuclear cells (PBMCs) from a healthy donor (4K PBMCs and 8K PBMCs) [29]. We derived quality control metrics using the cellrangerRkit R package (v. 1.1.0). Quality metrics were extracted from CellRanger throughout the molecule specific information file. After filtering as in [46], we extract 12,039 cells with 10,310 sampled genes and get biologically meaningful clusters with the software Seurat [47]. We then filter genes that we could not match with the bulk data used for differential expression to be left with *g* = 3346.

#### BRAIN LARGE

This dataset contains 1.3 million brain cells from 10x Genomics [20]. We randomly shuffle the data to get a 1M subset of cells and order genes by variance to retain first 10,000 and then 720 sampled variable genes. This dataset is then sampled multiple times in cells for the runtime and goodness-of-fit analysis. We report imputation scores on the 10k cells and 720 genes samples only.

#### RETINA

The dataset of bipolar cells from [30] contains after their original pipeline for filtering 27,499 cells and 13,166 genes coming from two batches. We use the cluster annotation from 15 cell-types from the author. We also extract their normalized data with Combat and use it for benchmarking.

#### HEMATO

This dataset with continuous gene expression variations from hematopoeitic progenitor cells [31] contains 4,016 cells and 7,397 genes. We removed the library *basal-bm1* which was of poor quality based on authors recommendation. We use their population balance analysis [48] result as a potential function for differentiation.

#### CBMC*

This dataset that includes 8,617 cord blood mononuclear cells [32] profiled using 10x along with for each cell 13 well-characterized mononuclear antibodies. We kept the top 600 genes by variance.

#### BRAIN SMALL*

This dataset consists in 9,128 mouse brain cells profiled using 10x [20] is used as a complement of PBMC for our study of zero abundance and quality control metrics correlation with our generative posterior parameters. We derived quality control metrics using the cellrangerRkit R package (v. 1.1.0). Quality metrics were extracted from CellRanger throughout the molecule specific information file. We kept the top 3000 genes by variance. We used the clusters provided by cellRanger for the correlation analysis of zero probabilities.

### 4.6 Algorithms used for benchmarking

#### Factor analysis

We used the factor analysis (FA) method from the scikitlearn python package. FA is always applied to log-data.

#### ZIFA

We used the zero-inflated factor analysis method (ZIFA) from https://github.com/epierson9/ZIFA with default parameters. We always apply ZIFA to log-data.

#### SIMLR

We used the large scale version of the Single-cell Interpretation via Multi-kernel LeaRning (SIMLR) algorithm from Bioconductor with parameters recommended by authors (k=30, kk=200). We always apply SIMLR to log-data as advocated in the original paper. We used the number of cluster as the number of cell-types. Except for the HEMATO data and the random ZINB dataset where we used the procedure SIMLR_Estimate_Number_of_Clusters.

#### BISCUIT

We applied the BISCUIT algorithm from https://github.com/sandhya212/BISCUIT_SingleCell_IMM_ICML_2016 with default parameters. As the code to recover the exact results from [10] was not available, we did not consider BISCUIT in the clustering benchmarking. Also, we checked different parameters for the number of iterations and the spin parameter without being able to get a configuration that would provide a better score for imputation.

#### ZINB-WaVE

We applied the ZINB-WaVE procedure from the R package zinbwave with the gene-level covariate to be a column of one and the cell-level covariate to be a column of ones. We always apply ZINB-WaVE to count-data.

#### PCA

We used the Principal Component Analysis method from the scikit-learn python package. We always apply PCA to log-data.

#### MAGIC

We used the Python3 code of the Markov Affinity-based Graph Imputation of Cells algorithm from https://github.com/KrishnaswamyLab/magic. We used the parameters from their iPython notebook on the github repo.

#### MAST

We used the R package MAST on log-counts to provide our differential expression analysis.

#### DESeq2

We used the R package DESeq2 on raw counts to provide our differential expression analysis.

#### edgeR

We used the R package edgeR on raw counts to provide our differential expression analysis.

#### ComBat

We used the R package sva on the bipolar dataset following the original preprocessing steps of the bipolar paper [30].

#### tSNE

We used the TSNE class from scikit-learn with default perplexity parameter of 30.

#### Force-directed Layout

We used the spring_layout module from the net-workx package by working with the raw 5-nearest neighbors adjacency matrix.

### 4.7 Evaluation

#### Log-likelihood on held-out data

We provide a multi-variate metric of goodness of fit on the data. The common method of evaluation for generative models is to evaluates the model on data it has never seen. Let us start with a complete dataset *X*. From this dataset, we randomly selected a training set *X_train_* and a held-out testing set *X_test_*. Each model gives and underlying probability measure *p* that we will specify. We define the log-likelihood on held-out data by integrating in the following way:

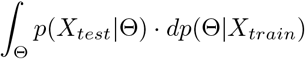

where *dp*(Θ|*X_train_*) designates the posterior parameters of the model after having fitted the training data and where *p*(*X_test_|*Θ) is assessing the goodness-of-fit of the held-out data under the chosen parameter Θ.

In the case of a fully generative model like BISCUIT, the posterior parameter *dp*(Θ| *X_train_*) is designated by a probability measure we have to sample from. For technical reasons, we did not include BISCUIT in this analysis. Specifically, the original code does not provide a straightforward way to evaluate posterior likelihood, on unseen data. In most generative models, we take a point estimate over these parameters (parameters of a neural network in scVI, factor loading matrix or decay rate in ZIFA) and the former measure is a Dirac centered on the parameters fixed at testing time.

Now, we focus on the value *p*(*X_test_*|Θ) itself. First, because some algorithms are run on log transformed data and some on raw data, we take into account the distortion into account for the log-likelihood scores (Appendix C). Second, this quantity can often be intractable because of latent variables we have to marginalize out. In that case, we can take lower bounds from the Expectation Maximization algorithm for ZIFA (Appendix E), exact value for FA and the variational lower bound for scVI (Appendix F). In the case of an algorithm where the latent variable are actual parameters to optimize as in ZINB-WaVE, we need to re-run this partial optimization at testing-time (Appendix D).

#### Corrupting the datasets for imputation benchmarking

We use in the paper two different approaches to measure the robustness of algorithms to noise in the data:

- Uniform zero introduction: We take randomly ten percent of the non-zero entries and multiply the entry *n* with a Ber(0.9) random variable.
- Binomial data corruption: We randomly select 10% of the matrix and replace an entry *n* by a Bin(*n*, 0.2) random variable.

#### Accuracy of imputing missing data

As imputation tantamount to replace missing data by its mean conditioned on being observed, we use the median 𝕃_1_ distance between the original dataset and the imputed values for corrupted entries only.

We now define what the imputed values are. For MAGIC, we use the output of their algorithm. For BISCUIT, we used the imputed counts. For ZIFA, we use the mean of the generative distribution conditioned on the non-zero event (mean of the factor analysis part) that we project back into count space. For scVI and ZINB-WaVE, we use the mean of the Negative Binomial distribution.

#### Silhouette width

The silhouette width requires either a similarity matrix or a latent space. We can define a silhouette score for each sample *i* with:

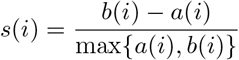

where *a*(*i*) is the average distance of *i* to all data points in the same cluster *c_i_*. *b*(*i*) is the lowest average distance of *i* to all data points in the same cluster *c* among all clusters *c*. Clusters can be replaced by batches if we are estimating the silhouette width for assessing batch effects [46].

#### Clustering metrics

The following metrics requires a clustering and not simply a similarity matrix. For these ones, we will use a K-means clustering on the given latent space with *T* = 200 random initializations to have a stable score.

#### Adjusted Rand Index

This index requires a clustering. Most

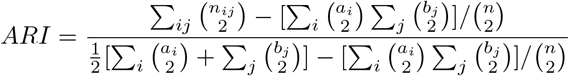

where *n_ij_, a_i_, b_j_* are values from the contingency table.

#### Normalized Mutual Information

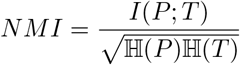

where *P, T* designates empirical categorical distributions for the predicted and real clustering. *I* is the mutual entropy and ℍ is the Shannon entropy.

#### Entropy of batch mixing

Fix a similarity matrix for the cells and take *U* to be a uniform random variable on the population of cells. Take *B_U_* the empirical frequencies for the 50 nearest neighbors of cell *U* being a in batch *b*. Report the entropy of this categorical variable and average over *T* = 100 values of *U*.

#### Protein abundance / mRNA expression

Take the similarity matrix for the normalized protein abundance (centered log-ratio transformation, see [32], Methods). Compute a 100 nearest neighbors graph. Fix a similarity matrix for the cells and compute a 100 nearest neighbors graph. Report the Spearman correlation of the flattened matrices and the fold enrichment.

Let *A* be the set of edges in the protein NN graph, *B* the set of edges in the cell NN graph and *C* the entire set of possible edges. The fold enrichment is defined as:

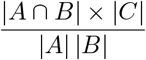

#### Baseline p-values from microarray data

For B cells against D cells, we used GSE29618) and between CD4+ and CD8+ T cells, we used GSE8835. We then used these reference gene sets to test the association of each gene’s expression with biological class, defining a two-sided t-test p-value per gene.

#### Differential Expression metrics

We used for each point of 100 cells from each cluster. In scVI, we draw 200 samples from the variational posterior. This subsampling ensures that our results are stable.

#### Area Under the Curve

We report the AUROC at detecting the most significantly expressed genes in the micro-array data (in all cases corrected p-values < 0.05).

#### Irreproducible Discovery Rate

The IDR is computed using the corresponding R package. We adjust the prior for the mixture weight to be the fraction of genes detected in the micro-array data.

## A Marginalizing out the latent variables of scVI

First, take *r* to be the gene-specific shape parameter of a Gamma variable *w*, 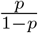 to be its scale parameter, and use a scalar λ ∈ ℝ^+^ then the count variable *y|w ~* Poisson(λ*w*) has a Negative Binomial marginal distribution with mean 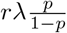

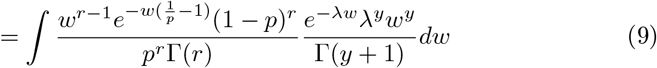

Second, multiplication by zero to *y_ng_* can be formally encoded as a mixture between a point-mass at zero and the original distribution of *y_ng_*.

Consequently, our conditional *p*(*x_ng_|z_n_, ℓ_n_, s_n_*) is a zero-inflated Negative Binomial with probability mass function:

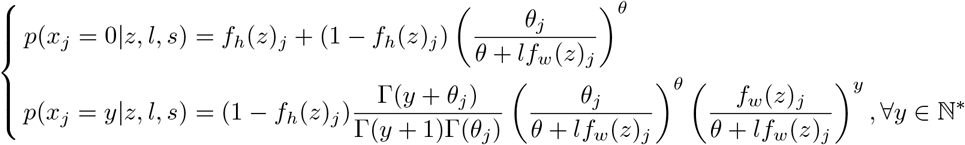

where *f_h_*(*z*) is encoding the zero probability of *h* and *f_w_*(*z*) the mean of *w*.

## B Negative binomial parametrization

**Negative binomial PMF parametrization** A choice of parametrization is crucial for optimization consideration. We could follow [9] by using a mean *μ* and an inverse-dispersion *θ* parameter:

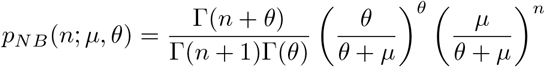

We also keep in mind a more gentle parametrization with nicer form even though still non-convex:

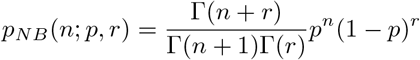

with 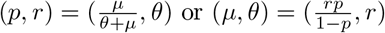

Because the first parametrization has a better behavior when scaling the Poisson mean as we do with library size normalization, this is the one we retain.

### Equivalence of parametrization

Assume now one wants to simulate what would have been the rate of the latent corresponding Poisson variable, one has to sample from a Gamma of shape 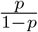 and scale 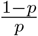

### Numerical considerations

We transformed the expression to incorporate logits and use Tensorflow numerically stable functions. Instead of writing explicitly a sigmoid non-linearity, the probability of zero in the mixture is given by:

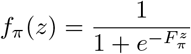

where 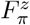 is the output of the neural network without non-linearity. We then write the log-likelihood as a function of 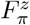.
*r* that can either be parametrized by a neural net or constant for each gene will be kept noted *r* for simplicity. 𝑆 denotes the softplus function *x* ⟼ log(1 + *e^x^*).

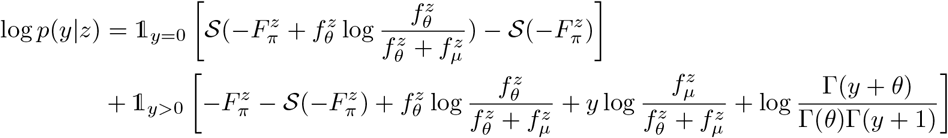

## C Comparing log-likelihood for log and non-log data

Let *X* be a positive random variable and let us note *Y* = log(1+*X*) and suppose we have a model for *Y* written ℙ_*Y*_. The likelihood score on the raw data is given by evaluating the density ℙ_*X*_ which is:

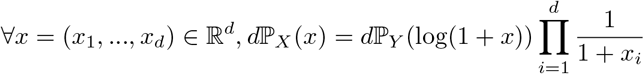

so this yield for the likelihood scores:

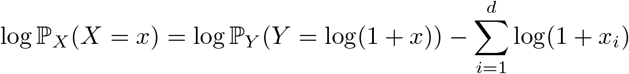

## D Log-likelihood for ZINB-WaVE

The function to be optimized for ZINB-WAVE is essentially penalized likelihood. One can thus run once the full optimization function on a training set as follow:

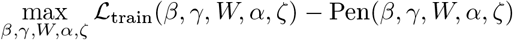

This optimization is performed by alternating minimization. By fixing the variables *β, α, ζ* learned from the training set, we can compute a likelihood on a validation set by performing inference over the latent variables *γ, W* which is a simple Ridge that can be solved in parallel by a simple modification of their code.

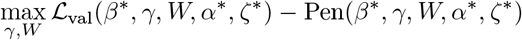

## E Log-likelihood for ZIFA

The EM algorithm naively provide a lower bound on the log-likelihood for ZIFA:

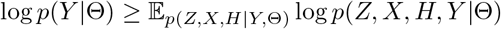

The complete log-likelihood has a simple expression:

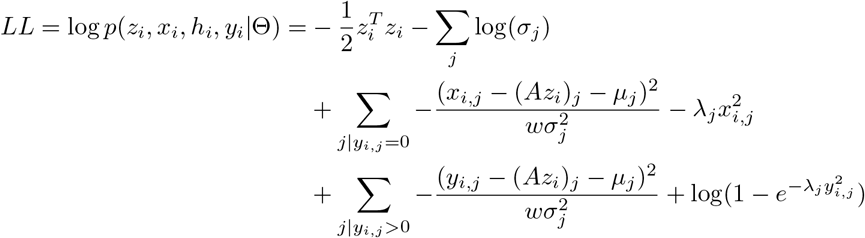

and the prior distribution is close to Gaussian so we can modify ZIFA code and use a E step to compute the desired value. E-step gives us the following values:

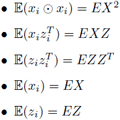

Then we have:

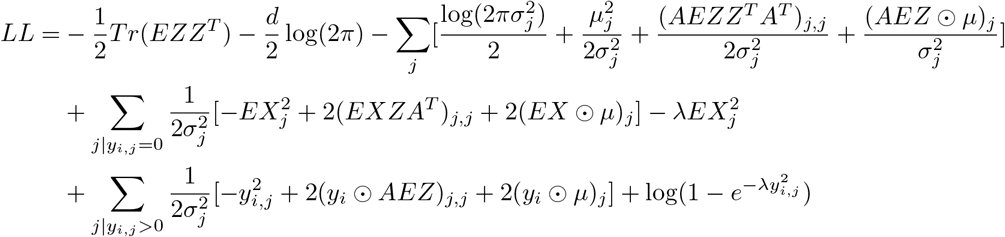

## F Log-likelihood for scVI

Our variational inference procedure provides us with a lower bound on the log-likelihood of held-out data:

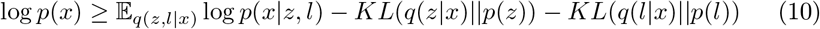

The lower-bound is tight whenever *q*(*z|x*) = *p*(*z|x*). Keeping the generative model as fitted on training data, we can optimize our inference network at test-time to have a better lower-bound of the held-out log-likelihood and report the best value. That is essentially equivalent to assess the marginal likelihood of held-out data, conditioned on a latent representation learned for the held-out data.

